# Reconstitution of +1 nucleosome transcription reveals coordinated functions of SAGA, Mediator, and TFIIH

**DOI:** 10.64898/2026.09.24.754195

**Authors:** Shigeki Nagai, Dong-Hua Chen, Rayees U.H. Mattoo, Heqiao Zhang, Roger D. Kornberg

## Abstract

The +1 nucleosome has emerged as a key regulator of eukaryotic transcription, but how it controls transcription initiation remains poorly understood. Here we reconstitute transcription through the +1 nucleosome using eleven purified yeast factors: RNA polymerase II (Pol II), the six general transcription factors (GTFs), TFIIS, the activator Pho4, and the SAGA and Mediator complexes. The system recapitulates key features of regulation observed in vivo. SAGA, acting with Pho4, directs pre-initiation complex (PIC) assembly to the correct position through its TBP-loading activity. Mediator stimulates transcription when the +1 nucleosome imposes a barrier to PIC formation, consistent with stabilization of productive TFIIH–DNA engagement. Contrary to the prevailing model, SAGA remains bound to the PIC after TBP loading and acetylates the +1 nucleosome within the assembled complex. The isolated PIC–Mediator–SAGA–nucleosome complex is transcriptionally active, and the repressive effect of the nucleosome is relieved by the DNA translocase activity of Ssl2, the TFIIH subunit that opens promoter DNA. TFIIH thus couples promoter melting to remodeling of the +1 nucleosome.

## INTRODUCTION

Genes of the yeast *Saccharomyces cerevisiae* fall into two classes: those that are constitutively expressed (“housekeeping” genes) and those that are inducible (repressed except when triggered by transcriptional activator proteins). All genes undergoing transcription exhibit a characteristic promoter chromatin architecture, with a nucleosome-free region of about 150 bp upstream that includes TBP-associated sequences (TATA-like or TATA-less at constitutively expressed promoters and a TATA box at inducible promoters), and with the transcription start site located within the first nucleosome downstream (the +1 nucleosome) (Chen and Pugh 2021). This architecture is the sole state of constitutively expressed promoters (Albert et al. 2007), whereas it arises at inducible promoters upon transcriptional activation (Nocetti and Whitehouse 2016).

Transcriptional activator proteins recruit four multiprotein complexes – SWI/SNF, SAGA, Mediator, and TFIID – both to create promoter chromatin architecture and to introduce TBP, which nucleates formation of a transcription pre-initiation complex (PIC). SWI/SNF is believed to remove the upstream nucleosome and to position the +1 nucleosome at inducible promoters (Nocetti and Whitehouse 2016; Kubik et al. 2019). SAGA contains an acetyltransferase responsible for acetylation of promoter nucleosomes and also delivers TBP to inducible promoters (Soffers and Workman 2020). TFIID brings TBP to most constitutively transcribed promoters (Chen and Pugh 2021). Finally, Mediator enters early and is required for transcription of all promoters, where it serves both as an integral component of the PIC and as the principal conduit of regulatory information to the transcription machinery (Richter et al. 2022).

Genetic studies have revealed overlapping roles of these multiprotein complexes. SAGA contributes to transcription at constitutively transcribed promoters, likely through its histone H3 acetylation and histone H2B deubiquitylation activities (Baptista et al. 2017; Donczew et al. 2020; Mittal et al. 2022), and TFIID acts at some inducible promoters (Warfield et al. 2017; Donczew et al. 2020; Mittal et al. 2022). Biochemical studies are needed to disentangle the roles of these complexes and to define their molecular mechanisms. The +1 nucleosome is of particular interest, and indeed presents a paradox: it is required for transcription, and yet a nucleosome covering a transcription start site is expected to prevent initiation. We have therefore performed biochemical studies in a fully defined transcription system with DNA templates bearing positioned nucleosomes. The results give insight into the coordinated functions of SAGA, Mediator, and TFIIH in transcription through the +1 nucleosome.

## RESULTS

### Reconstitution of +1 nucleosome transcription at inducible promoters

We developed an experimental system that allows site-specific installation of a mononucleosome downstream of native *Saccharomyces cerevisiae* TATA-containing, stress-induced promoters, including *PHO5*, *HIS4*, and *TEA1*. A 160 bp DNA fragment containing the transcription start sites (TSSs) and part of the 601 positioning sequence was digested with the type IIS restriction enzyme BsaI, which cleaves outside its recognition sequence, and was assembled into a mononucleosome by salt-gradient dialysis followed by sucrose gradient centrifugation. The purified mononucleosome was ligated to BsaI-digested promoter DNA and immobilized on magnetic beads through a biotin tag at the upstream end of the promoter (Supplemental Fig. S1A). Naked DNA templates were prepared in parallel by treating the same nucleosomal templates with 2 M NaCl to strip histones from the DNA, so that nucleosomal and naked templates differ only in the presence of histones. Restriction endonuclease digestion confirmed correct positioning of the nucleosome (Supplemental Fig. S1B).

The +1 nucleosome and the corresponding naked DNA templates were transcribed with defined factors, including Pol II, the six GTFs, TFIIS, the Pho4 activator, Mediator, and SAGA (Supplemental Fig. S2). The transcription assay was performed as follows (Figure 1A): all components were first added to the immobilized template to assemble the PIC in the absence of ATP; unbound factors were removed by washing; and NTPs were then added to initiate transcription. Transcripts were labeled by incorporation of [α-^32P^]-UTP, allowing both specific and nonspecific products to be visualized. Because PIC assembly precedes the addition of NTPs and is followed by a wash, the assay reports on the formation and retention of transcription-competent complexes. We initially installed the +1 nucleosome 30 bp downstream of the 5′-edge of the TATA box of the *PHO5* promoter (T30N), a distance associated with maximal *PHO5* expression *in vivo* (Small et al. 2014).

**Figure 1.**
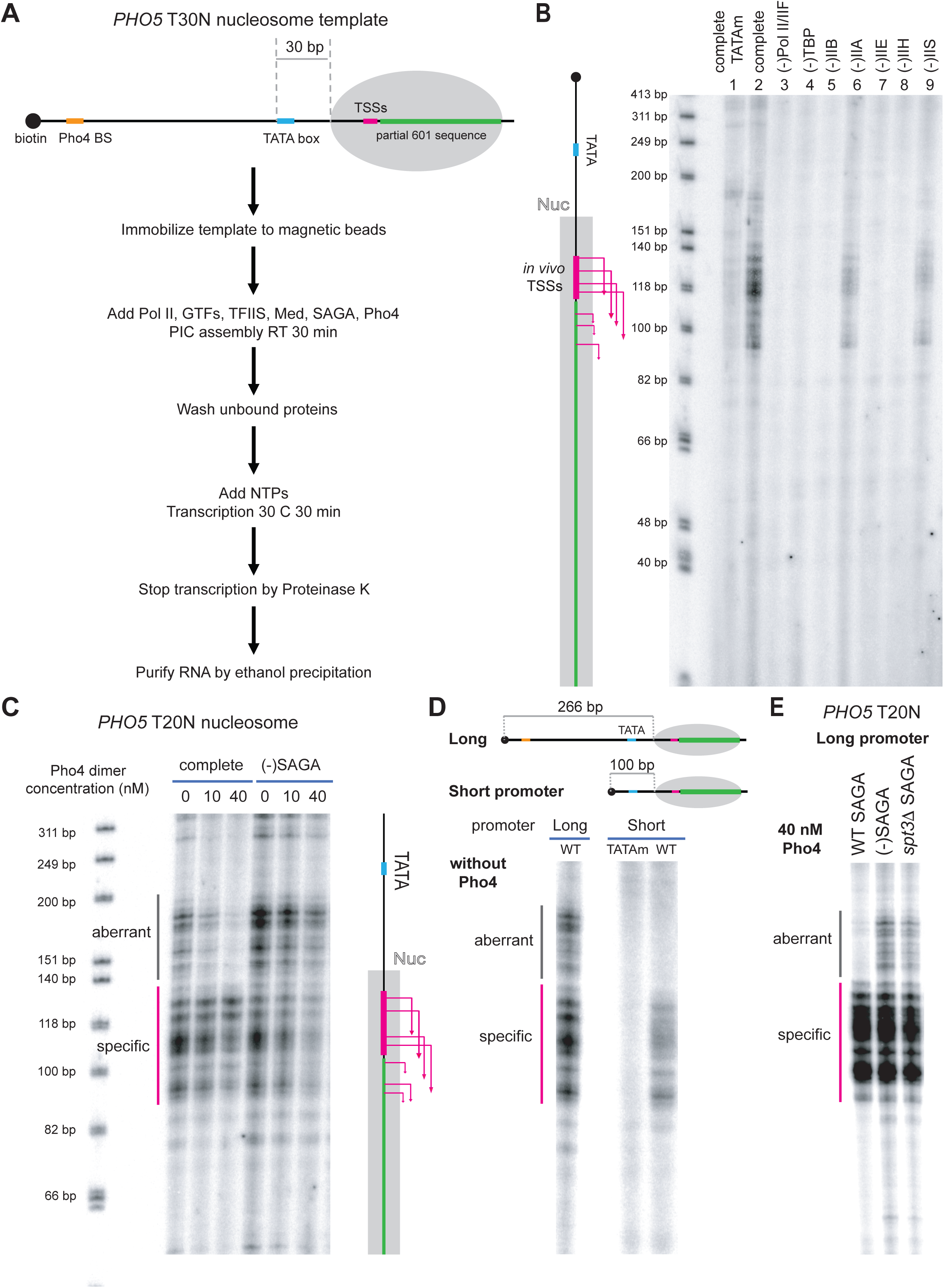
Immobilized +1 nucleosome transcription assay. (A) Scheme of the immobilized +1 nucleosome transcription assay. The +1 nucleosome was installed 30 bp downstream of the 5′-edge of the TATA box of the *PHO5* promoter (T30N). (B) Transcription reactions performed as in (A) on the *PHO5* T30N nucleosome template. The complete reaction is shown in lane 2; lane 1 used a template with a mutated TATA box (TATAm); the remaining reactions contained all factors except the component indicated above each lane. (C) Pho4 titration. Transcription reactions were performed as in (A) with the *PHO5* T20N nucleosome template, with the complete set of factors or in the absence of SAGA. Pho4 represses aberrant transcripts, and this effect is lost in the absence of SAGA. (D) Aberrant transcripts arise from the region 80–246 bp upstream of the *PHO5* TATA box. Transcription reactions without Pho4 were performed on the *PHO5* T20N nucleosome with either the long or the short promoter; the short promoter lacks this upstream region. (E) Substituting *spt3*Δ SAGA for wild-type SAGA resulted in numerous aberrant transcripts. Reactions were performed on the long *PHO5* T20N promoter template in the presence of 40 nM Pho4.

Transcription of the *PHO5* +1 nucleosome with the complete set of factors generated TATA-dependent transcripts, as evidenced by their absence from a template with a mutated TATA box (Figure 1B, compare lanes 1 and 2). Most transcripts initiated within the previously described 20 bp initiation region located 50–70 bp downstream of the TATA box (Figure 1B, magenta lines in the cartoon) (Nagai et al. 2017). Transcripts shorter than the predicted sizes were also observed (Figure 1B, lane 2), and these too required the TATA box (Figure 1B, compare lanes 1 and 2). The shorter transcripts likely initiate from downstream sites within the 601 sequence through TSS scanning, a process driven by the DNA translocase activity of TFIIH (Fishburn et al. 2015; Murakami et al. 2015a). Pol II and five GTFs (TBP, TFIIB, -IIE, -IIF, and -IIH) were absolutely required for transcription, and TFIIA and TFIIS strongly stimulated transcription of the +1 nucleosome (Figure 1B, lanes 6 and 9). We refer to these TATA-dependent transcripts hereafter as “specific” transcripts.

### Recruitment of SAGA by Pho4 represses aberrant transcripts on the +1 nucleosome

On the *PHO5* T20N nucleosome template, omission of Pho4 resulted in numerous aberrant transcripts (Figure 1C). We reasoned that suppression of aberrant transcripts is mediated through Mediator or SAGA, both of which are known to interact with activator proteins. SAGA is recruited to the *PHO5* and *PHO8* promoters by Pho4 *in vivo* (Barbaric et al. 2003; Biddick and Young 2009). Omission of Mediator reduced specific transcripts but did not give rise to aberrant transcripts (Figure 2B). In contrast, omission of SAGA resulted in numerous aberrant transcripts even in the presence of Pho4 (Figure 1C). SAGA can efficiently deliver TBP to DNA containing a canonical TATA box or a TATA-like element with one or two mismatches, but not to DNA lacking any such element (Papai et al. 2020). SAGA evidently prevents delivery of TBP to spurious DNA sequences, thereby directing PIC assembly to the correct location.

**Figure 2.**
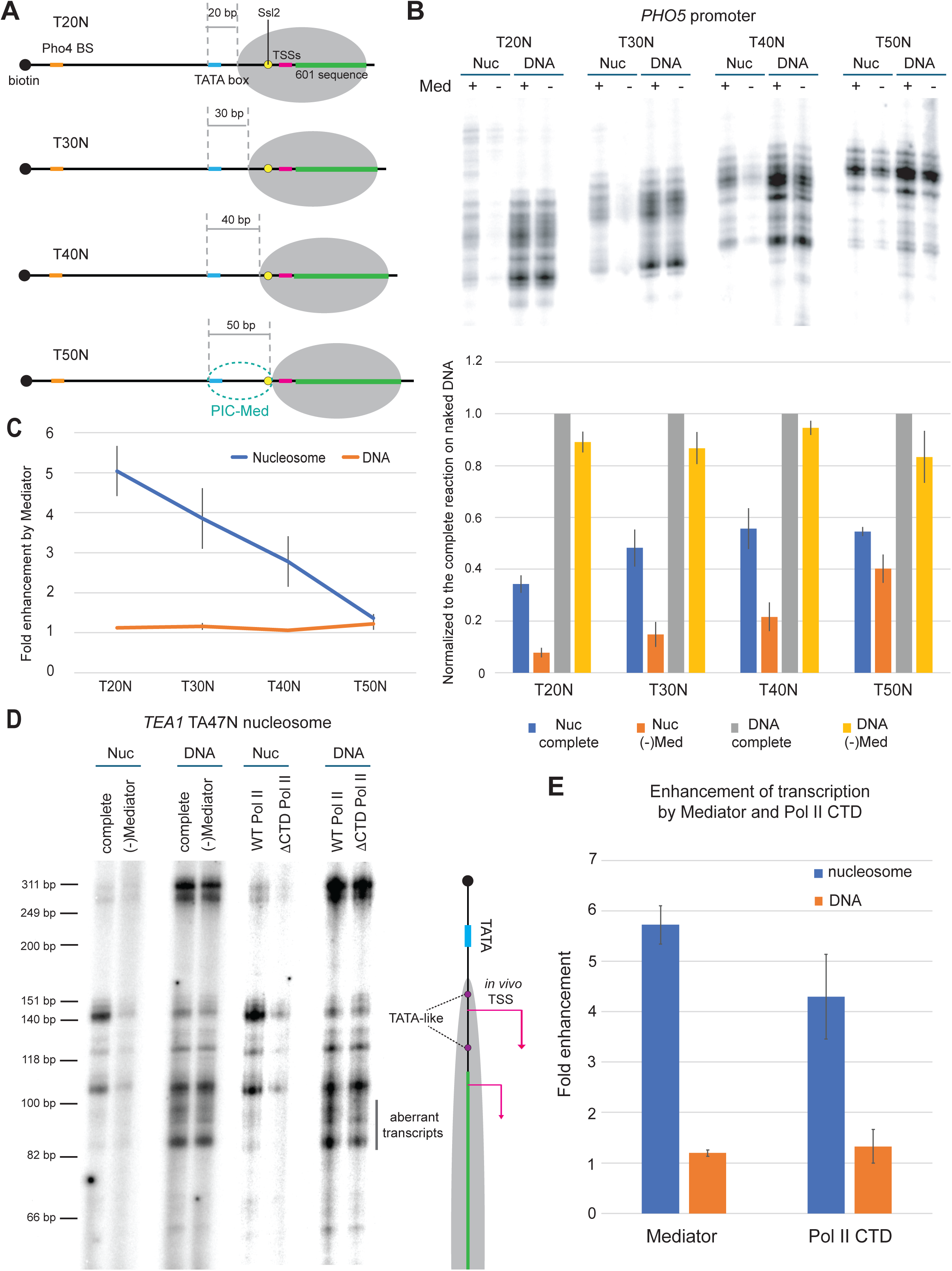
Mediator dependence varies with the position of the +1 nucleosome. (A) The +1 nucleosome was installed 20, 30, 40, or 50 bp downstream of the 5′-edge of the *PHO5* TATA box. The region of promoter DNA occupied by PIC-Med is indicated by the dashed green circle. (B) (Top) Transcription was performed with the indicated configurations of the +1 nucleosome and the corresponding naked DNA templates, with or without Mediator. (Bottom) Quantitation of specific, promoter-dependent transcription. For each configuration, the signal was normalized to that from naked DNA with the complete set of factors (grey bars). Values are means of 3–6 independent experiments ± SEM. (C) Fold enhancement of transcription by Mediator. For each nucleosome or naked DNA template, specific transcription signal from the complete reaction was divided by the signal from the reaction without Mediator. Values are mean of 3–6 independent experiments ± SEM. (D) Transcription of the inducible *TEA1* promoter with an associated +1 nucleosome (T47N). The complete reaction was performed as in (B). Other reactions contained all factors but lacked Mediator, or contained ΔCTD Pol II in place of wild-type Pol II. Two TATA-like elements covered by the +1 nucleosome are indicated by purple dots (cartoon, right); the aberrant transcripts from the naked DNA template likely arise from these elements. (E) Fold enhancement of transcription by Mediator or by the Pol II CTD. For each nucleosome or naked DNA template, the specific transcription signal from the complete reaction was divided by that from the reaction lacking Mediator or from the reaction with ΔCTD Pol II. Values are means of 3 independent experiments ± SEM.

The aberrant transcripts arise from a region 80–246 bp upstream of the TATA box, since deleting this region eliminated them in the absence of Pho4 (Figure 1D). A PIC therefore appears to form at a spurious sequence within this upstream region. Suppression of aberrant transcripts by SAGA is largely mediated by its Spt3 subunit, which loads TBP onto TATA and TATA-like elements (Papai et al. 2020): substituting *spt3*Δ SAGA for wild-type SAGA resulted in numerous aberrant transcripts (Figure 1E; Supplemental Fig. S2).

SAGA is also required to suppress aberrant transcription on naked DNA, but this suppression does not require Pho4 (Supplemental Fig. S3A). The proximal +1 nucleosome therefore appears to create a requirement for activator-assisted SAGA function when it impedes productive PIC assembly. Under these conditions, Pho4 may recruit or stabilize SAGA at the intended promoter, favoring selective TBP delivery and limiting inappropriate initiation at accessible upstream sites, whereas on naked DNA SAGA maintains initiation specificity without this additional contribution from Pho4. Simultaneous interactions of SAGA with Pho4 and with the +1 nucleosome may further constrain its positioning, favoring TBP delivery to the promoter TATA element (Figure 6, step 2). The precise mechanism remains to be established, but these findings support a role for Pho4 in maintaining SAGA-dependent initiation specificity in the presence of a nucleosomal barrier.

### Mediator dependence increases with promoter proximity of the +1 nucleosome

The dependence of transcription on Mediator varied with the position of the +1 nucleosome, whereas Mediator had little effect on the corresponding naked DNA templates (Figure 2). Transcription was tested with the +1 nucleosome installed 20, 30, 40, or 50 bp downstream of the 5′-edge of the TATA box (Figure 2A). The +1 nucleosome repressed transcription by the minimal set of components (Pol II, GTFs, and TFIIS), and repression was stronger the more proximally the nucleosome was positioned (Figure 2B, orange bars), consistent with a previous study (Abril-Garrido et al. 2023). Importantly, this repression was largely relieved by Mediator, which had minimal effect on transcription of naked DNA (Figures 2B and 2C). The effect of Mediator was more pronounced the more proximally the +1 nucleosome was positioned (Figure 2C): Mediator enhanced transcription approximately 5-fold on the T20N template but only ∼1.3-fold on the T50N template. A similarly strong Mediator dependence was observed for transcription of the *HIS4* T27N +1 nucleosome, but not of the corresponding naked DNA (Supplemental Fig. S3B).

A previous genome-wide study showed that the +1 nucleosome lies on average 15–20 bp from the 5′-edge of the TATA box (or TATA-like sequence) at actively transcribed yeast genes (Nocetti and Whitehouse 2016). Mediator is strongly required for transcription in this configuration *in vitro* (Figure 2C). Mediator-dependent transcription was also observed in the absence of SAGA (Supplemental Fig. S3C), indicating that this dependence does not require SAGA. Together, these results indicate that Mediator becomes particularly important when the position of the +1 nucleosome constrains productive PIC assembly.

Assembly of the PIC on the immobilized *PHO5* T30N nucleosome showed that TBP is efficiently recruited to the promoter independently of Mediator, whereas Pol II recruitment is strongly enhanced by Mediator (Supplemental Fig. S4). Mediator thus promotes PIC assembly at a promoter where a proximally positioned +1 nucleosome imposes a barrier to PIC formation.

### A model for stabilization of a productive PIC by Mediator

A cryo-EM structure of the human +1 nucleosome-bound PIC-Mediator (Med)-TFIID showed that Mediator contacts the +1 nucleosome through its hook domain, and the authors proposed that this interaction contributes to transcription of the +1 nucleosome (Chen et al. 2022). In contrast, a cryo-EM structure of the yeast +1 nucleosome-bound PIC-cMed (core Mediator: Mediator lacking the Tail module) showed no direct contact between Mediator and the nucleosome (Schilbach et al. 2023). Our finding that Mediator dependence increases as the nucleosome approaches the promoter suggests that Mediator facilitates transcription by stabilizing a productive PIC under conditions in which nucleosomal constraints oppose its assembly, rather than by contacting the nucleosome directly.

In the *PHO5* position series, Mediator dependence was strongest when the +1 nucleosome was installed 20 bp from the 5′-edge of the TATA box and declined as the nucleosome was moved to 50 bp (Figures 2B and 2C). Cryo-EM analyses of yeast and human PIC-Med have placed the Ssl2 subunit of TFIIH (XPB in human) at the downstream edge of the PIC, engaging promoter DNA approximately 45–50 bp from the 5′-edge of the TATA box (Murakami et al. 2015b; He et al. 2016). A nucleosome positioned within this region can therefore interfere with productive TFIIH engagement. Consistent with this interpretation, the structure of a human PIC assembled on a T48N nucleosome showed partial unwrapping of the promoter-proximal nucleosomal DNA, allowing XPB to engage the promoter (Abril-Garrido et al. 2023).

Nucleosomal DNA transiently unwraps and rewraps (Li et al. 2005), and spontaneous unwrapping becomes less frequent at positions deeper within the nucleosome (Tims et al. 2011). Productive PIC assembly next to a proximal +1 nucleosome may therefore require capture and stabilization of a partially unwrapped state. The decrease in transcription as the nucleosome approaches the promoter (Figure 2B) is consistent with this constraint. We propose that Mediator provides additional stabilizing interactions within the PIC, increasing the probability that transiently exposed DNA is engaged productively and that the resulting complex persists. Such a mechanism requires neither direct contact between Mediator and the nucleosome nor an increase in the intrinsic rate of nucleosome unwrapping.

Previous cryo-EM analyses revealed substantial mobility of TFIIH within the PIC, with full engagement of TFIIH with TFIIE and promoter DNA observed in only a subset of particles (Murakami et al. 2015b; He et al. 2016; Tsai et al. 2017). When the +1 nucleosome is positioned close to the promoter, transient disengagement of TFIIH could allow partially unwrapped nucleosomal DNA to rewrap, restricting access of Ssl2 to downstream DNA and destabilizing the transcription-competent PIC. Stabilization of productive TFIIH–DNA engagement may therefore be particularly important when the +1 nucleosome constrains the incorporation of Ssl2 onto the promoter (Figure 6, step 4).

Structural studies of yeast PIC-Med have identified an interaction between the Mediator hook and the TFIIH subunit Rad3 (Robinson et al. 2016; Schilbach et al. 2017), providing a possible molecular basis for such stabilization. In our cryo-EM analysis of yeast PIC-Med, 3D variability analysis in cryoSPARC (Punjani and Fleet 2021) revealed coordinated motions of Mediator and core TFIIH (Supplemental Video 1), consistent with structural coupling between the two complexes, likely mediated by the hook-Rad3 interaction. These observations support a model in which Mediator stabilizes a transcription-competent PIC, potentially by favoring a TFIIH configuration that permits productive engagement of Ssl2 with downstream DNA. The specific contribution of the hook–Rad3 interaction to transcription of the +1 nucleosome remains to be established.

### Pol II CTD is crucial for transcription of the +1 nucleosome

We next examined transcription of the +1 nucleosome at the inducible *TEA1* promoter, which contains a TATA-like element (TATAAACG) differing by one nucleotide from the canonical TATA sequence. Nocetti et al. mapped the +1 nucleosome under different growth conditions and reported maximal transcription when the nucleosome was positioned approximately 47 bp downstream of the 5′-edge of the TATA element (Nocetti and Whitehouse 2016). Mediator enhanced transcription of the *TEA1* T47N nucleosome template approximately 6-fold but had little effect on the corresponding naked DNA template (Figures 2D and 2E). This stimulation exceeded the ∼2.6-fold enhancement observed on the *PHO5* T40N template, which contains a canonical TATA element. A noncanonical TATA element may thus increase the dependence on Mediator when the +1 nucleosome constrains PIC assembly, although other differences in promoter context may also contribute.

Transcription of naked DNA also yielded additional transcripts that were absent from the nucleosomal template (Figure 2D). These transcripts likely originate from two TATA-like elements within the region occupied by the +1 nucleosome (Figure 2D, cartoon, purple dots). Their persistence in the presence of SAGA is consistent with its ability to load TBP onto TATA-like elements, whereas their suppression on the nucleosomal template suggests that the +1 nucleosome restricts access to these alternative initiation sites. These results demonstrate suppression of aberrant transcription by the +1 nucleosome *in vitro*, consistent with its reported role *in vivo* (Kubik et al. 2019).

The carboxyl-terminal domain (CTD) of Pol II mediates its interaction with Mediator *in vitro* and is thought to contribute to assembly of the Mediator-associated PIC (Kim et al. 1994; Sogaard and Svejstrup 2007; Robinson et al. 2016). To examine its contribution to nucleosomal transcription, we prepared Pol II lacking the CTD (ΔCTD Pol II) from a yeast strain in which Protein A domains and a TEV protease cleavage site were inserted immediately upstream of the Rpb1 CTD repeats (Supplemental Fig. S2). Removal of the CTD strongly reduced transcription of the *TEA1* T47N nucleosome template but had little effect on transcription of naked DNA (Figures 2D and 2E). The parallel requirements for the CTD and for Mediator are consistent with a role for the Mediator–Pol II association in the formation or maintenance of a transcription-competent PIC on a nucleosomal template. Both requirements persisted in the absence of SAGA (Supplemental Fig. S3C).

We also examined the effect of the Mediator kinase module, which has been implicated in transcriptional repression *in vivo* (Luyties and Taatjes 2022). The four-subunit kinase module was purified from a yeast strain carrying Med12-TAP (Supplemental Fig. S3D). Addition of the purified Mediator kinase module strongly inhibited Mediator-dependent transcription of the +1 nucleosome (Supplemental Fig. S3D). The reconstituted system evidently recapitulates an inhibitory effect of the kinase module consistent with its reported repressive function *in vivo*.

### Isolation of a SAGA-bound PIC

Although SAGA functions in transcription initiation through TBP loading and acetylation of the +1 nucleosome (Soffers and Workman 2020; Mittal et al. 2022), it has remained unknown whether SAGA binds the PIC. Because SAGA binds both nucleosomes and activators, we first examined SAGA-PIC binding in the absence of a nucleosome and an activator. For this purpose we used the *SNR20* promoter, at which transcription initiates from two *in vivo* initiation sites without aberrant transcripts *in vitro* in the absence of an activator (Murakami et al. 2015a). SAGA was included in a procedure developed for assembly of the yeast PIC-Med (Robinson et al. 2016), in which Pol II, the six GTFs (TBP, TFIIA, -IIB, -IIE, -IIF, and -IIH), TFIIS, and SAGA are combined in a high-ionic-strength solution, dialyzed to physiological conditions, and isolated by 10%–40% glycerol gradient sedimentation (Figure 3A). Peak fractions contained all 50 PIC and SAGA polypeptides co-migrating at the same position in the gradient (Figures 3B and 3C) and were transcriptionally active *in vitro* (Figure 3D). Dynamic light scattering (DLS) of the isolated complex gave a cumulant hydrodynamic radius (Rh) of 21.82 ± 0.1 nm with a polydispersity index (PDI) of 0.06, indicating a highly monodisperse species, and substantially larger than the Rh of the PIC alone (14.31 ± 0.11 nm) (Figure 3E). The narrow, single-peak size distribution indicates that PIC-SAGA behaves in solution as a homogeneous species rather than a heterogeneous mixture of variably associated particles, consistent with specific incorporation of SAGA into the PIC.

**Figure 3.**
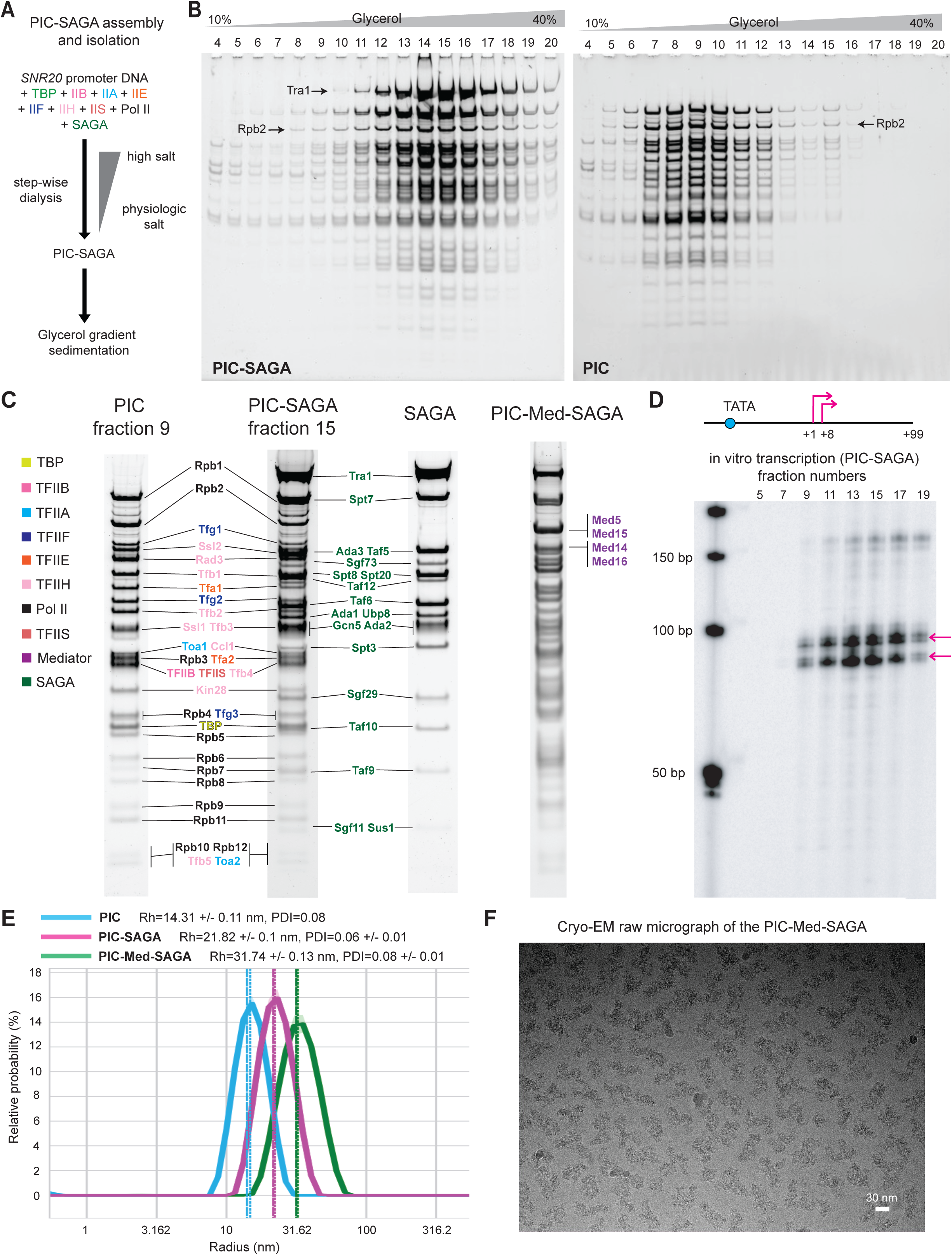
Isolation and characterization of a PIC-SAGA complex assembled on *SNR20* promoter DNA. (A) PIC-SAGA was assembled on *SNR20* promoter DNA by step-wise dialysis from high to physiological salt and isolated by 10%–40% glycerol gradient sedimentation. (B) SDS-PAGE of gradient fractions of yeast PIC-SAGA (left) or PIC (right). (C) SDS-PAGE of PIC (fraction 9), PIC-SAGA (fraction 15), SAGA alone, and PIC-Med-SAGA. (D) Transcripts from gradient fractions of PIC-SAGA, visualized by incorporation of [α-32P]-UTP and analyzed by gel electrophoresis and autoradiography. The *SNR20* promoter scheme used for PIC-SAGA assembly is shown above; transcription start sites (TSSs) are indicated by magenta arrows. (E) Dynamic light scattering (DLS) of PIC (light blue), PIC-SAGA (magenta), and PIC-Med-SAGA (green) without crosslinker. Data shown is the mean % relative probability distribution of hydrodynamic radius (*n* = 80 for each sample), with error values represented by the lighter color shade. (F) Representative raw micrograph of PIC-Med-SAGA assembled on the *SNR20* promoter DNA (scale bar: 30 nm).

Upon incorporation of Mediator into PIC-SAGA (Supplemental Fig. S5), we observed a further increase in complex size (cumulant Rh = 31.74 ± 0.13 nm, PDI = 0.08) (Figure 3E). PIC-Med-SAGA was subjected to cryo-EM analysis, for which glutaraldehyde crosslinking was performed by the gradient fixation method (GraFix) (Stark 2010). A cryo-EM data set revealed fields of monodisperse particles with an approximate diameter of 60 nm (Figure 3F), in good agreement with the hydrodynamic radius obtained by DLS in solution (cumulant Rh = 31.74 nm) (Figure 3E).

### Cryo-EM analysis of SAGA-containing PIC complexes

We next sought structural evidence for incorporation of SAGA into PIC-cMed by cryo-EM, comparing PIC-cMed-SAGA with a PIC-cMed control. Both complexes were assembled on the same minimal promoter fragment, extending 10 bp upstream and 52 bp downstream of the 5′-edge of the TATA box (Supplemental Fig. S6A), which is too short to accommodate SAGA bound elsewhere on the DNA and thereby rules out occupancy of a distal DNA site. In both maps, TBP, TFIIA, TFIIB, and TFIIH/Ssl2 were correctly positioned, confirming a properly assembled PIC (Supplemental Fig. S6B). Addition of SAGA produced additional density contacting TFIIH and TFIIS that was absent from the SAGA-free control; this density was too poorly resolved to be assigned to specific SAGA subunits (Supplemental Fig. S6B).

Although the resolution of the additional density does not allow the contributing subunits to be identified, its position, contacting TFIIH and TFIIS, agrees with previous reports of direct SAGA–TFIIH and SAGA–TFIIS contacts in yeast and human cells (Spt8–TFIIS: (Wery et al. 2004); Gcn5/KAT2A–Ssl2/XPB: (Sandoz et al. 2019)). These results support genuine incorporation of SAGA into the PIC, possibly in a conformationally dynamic configuration.

### Isolation of a SAGA-bound PIC assembled on the +1 nucleosome

Having established the SAGA-PIC interaction, we tested assembly of PIC-Med-SAGA on the +1 nucleosome, the physiological substrate for transcription initiation. At most transcriptionally active promoters, the TSS lies 10–15 bp inside the upstream border of the +1 nucleosome (Albert et al. 2007). To mimic this arrangement, we designed an *SNR20* promoter construct carrying a 601 positioning sequence such that the first TSS lies 15 bp inside the upstream border of the nucleosome (Figure 4A: *SNR20* T77N). PIC-Med-SAGA was assembled on the *SNR20* T77N nucleosome template and isolated by 10%–40% glycerol gradient sedimentation (Figure 4B). DLS of the isolated complex, without crosslinking, showed that it was highly monodisperse (PDI = 0.07 ± 0.01), with a cumulant Rh of 33.74 ± 0.13 nm (Figure 4C). Addition of acetyl-CoA to the isolated complex resulted in acetylation of histone H3 (Figure 4D), demonstrating engagement of the SAGA HAT module with the +1 nucleosome within the PIC. This is consistent with a previous report of SAGA-mediated acetylation of the +1 nucleosome *in vivo* (Mittal et al. 2022).

**Figure 4.**
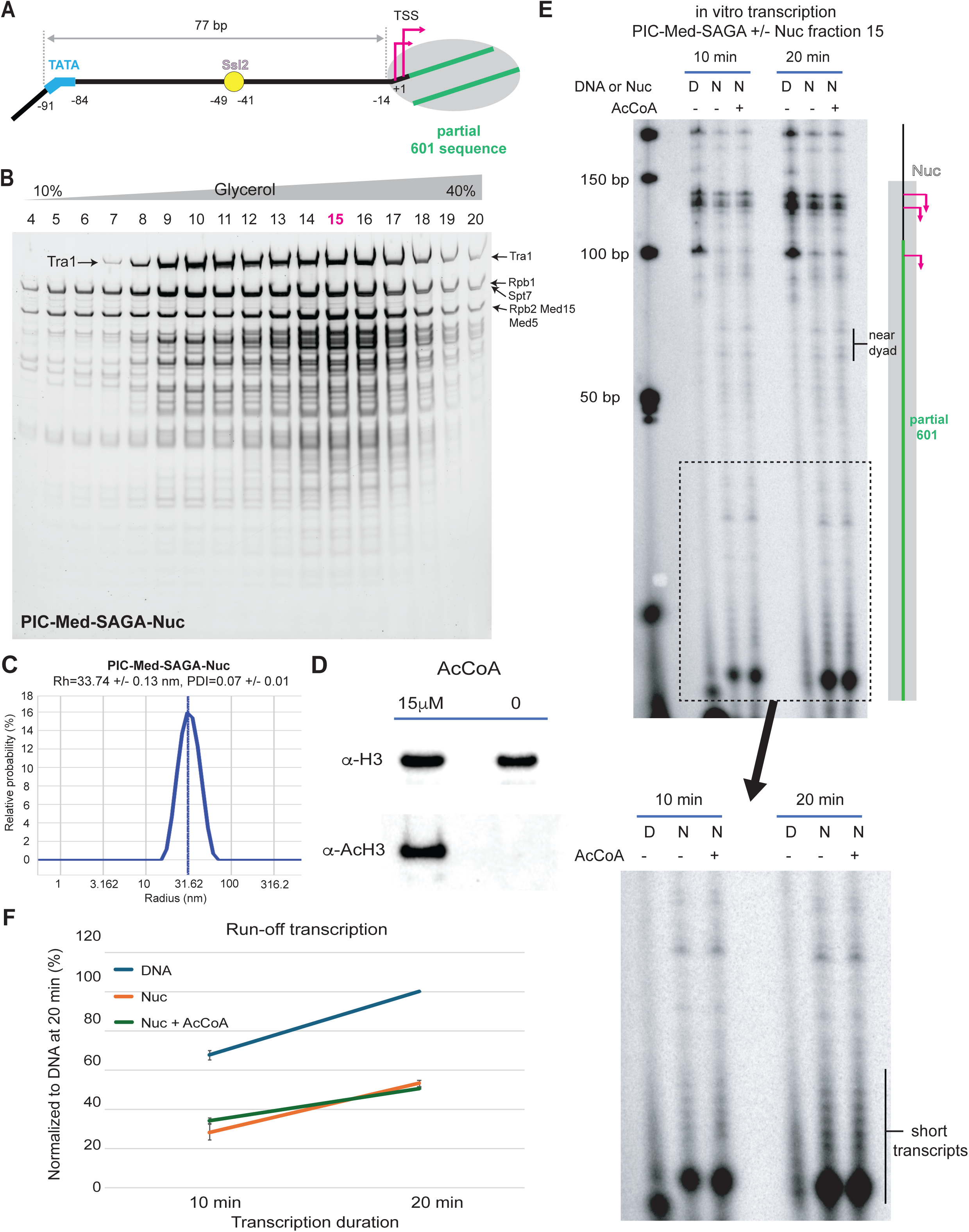
Isolation and characterization of a PIC-Med-SAGA-nucleosome complex assembled on *SNR20* promoter DNA. (A) Scheme of the +1 nucleosome-containing *SNR20* promoter (T77N) used for PIC-Med-SAGA-nucleosome assembly. Numbers indicate positions relative to the first TSS (+1). (B) PIC-Med-SAGA-nucleosome was assembled on the *SNR20* promoter with a positioned +1 nucleosome (Figure 4A) and isolated by 10%–40% glycerol gradient sedimentation. SDS-PAGE of gradient fractions is shown. (C) DLS of PIC-Med-SAGA-nucleosome. Data shown is the mean % relative probability distribution of hydrodynamic radius (*n* = 80 for each sample), with error values represented by the lighter color shade. (D) The isolated PIC-Med-SAGA-nucleosome complex (fraction 15) was treated with or without 15 µM acetyl-CoA for 30 min at room temperature. Acetylated histone H3 was detected by immunoblot using an antibody against pan-acetylated H3. (E) Time course of run-off transcription. Transcripts from gradient fraction 15 of PIC-Med-SAGA assembled on naked DNA (D) or on DNA with a +1 nucleosome (N) were labeled by incorporation of [α-32P]-UTP and analyzed by gel electrophoresis and autoradiography. Reactions on the nucleosome template were performed with or without 15 µM acetyl-CoA. An expanded view of the short-transcript region is shown below. (F) Quantitation of full-length transcripts. The specific transcription signal at each time point was normalized to that from the naked DNA template at 20 min. Values are means of 3 independent experiments ± SEM.

The isolated complex was transcriptionally active *in vitro*, with run-off transcription reduced by the presence of the nucleosome to ∼53% of the level obtained on naked DNA at 20 min (Figures 4E and 4F), consistent with the immobilized template assay (Figure 2B). This reduction reflects stalling of Pol II, including at sites near the dyad (Figure 4E), as shown previously for Pol II elongating through a nucleosome (Kireeva et al. 2005; Kujirai et al. 2018). Transcription of the +1 nucleosome template also gave rise to short (3–10 nt) RNA species (Figure 4E, lower enlarged panel), likely abortive or stalled products of early elongation. Addition of acetyl-CoA did not increase transcript levels (Figures 4E and 4F). Acetylation of the +1 nucleosome by SAGA may instead serve to recruit bromodomain-containing proteins such as RSC, SWI/SNF, and Taf14 (Hassan et al. 2002; Shanle et al. 2015; Lorch et al. 2018).

### TFIIH remodels the +1 nucleosome

Despite the premature transcripts arising from Pol II stalling, a significant fraction of the PIC-Med-SAGA-nucleosome complexes produced full-length transcripts (Figures 4E and 4F), suggesting that a component of the PIC relieves the repressive effect of the +1 nucleosome. Recent cryo-EM studies of the +1 nucleosome-bound PIC in yeast and human cells showed that TFIIH contacts the +1 nucleosome, with the Ssl2 translocase engaged with promoter DNA adjacent to the nucleosome (Chen et al. 2022; Abril-Garrido et al. 2023; Wang et al. 2023; Zhan et al. 2026). We therefore asked whether the +1 nucleosome is remodeled by the Ssl2 DNA translocase upon transcription initiation.

To test this, PIC-Med-SAGA was assembled on the immobilized *PHO5* T50N +1 nucleosome and treated with ATP, and the accessibility of the BsiWI and StyI sites within the nucleosome was assayed. The ClaI site, located upstream of the promoter, was used to release the template after the remodeling reaction. Whereas the BsiWI and StyI sites in naked DNA were almost completely accessible, both were strongly protected in the presence of the +1 nucleosome (Figure 5A). Upon addition of ATP, these sites within the nucleosome became accessible (Figure 5A). Remodeling was abolished when TFIIH was omitted from the reaction (Figure 5B), showing that TFIIH is responsible for remodeling of the +1 nucleosome.

**Figure 5.**
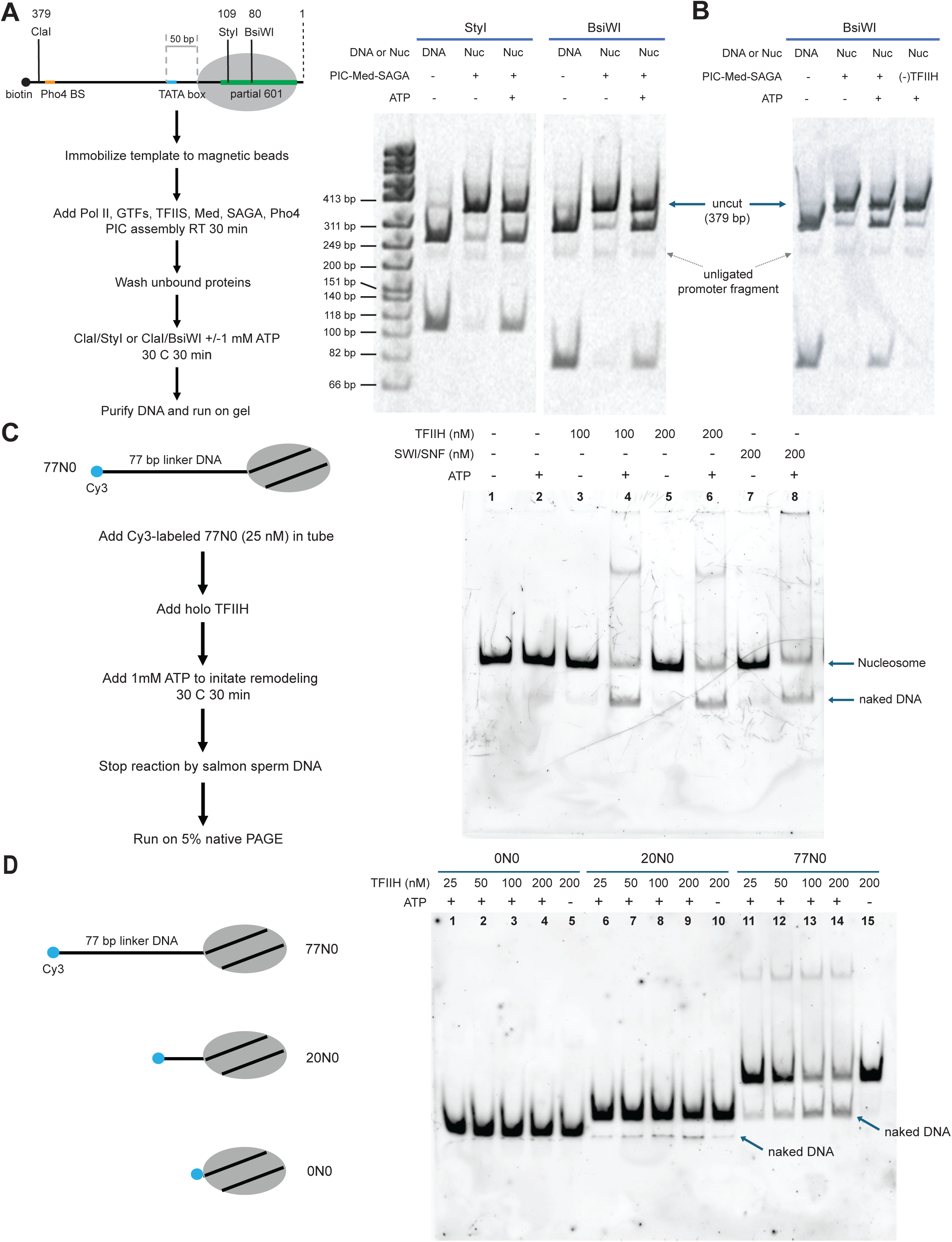
TFIIH remodels the +1 nucleosome. (A) Scheme of the immobilized +1 nucleosome remodeling assay with the *PHO5* T50N nucleosome template. PIC-Med-SAGA was assembled as in Figure 1A and incubated with or without 1 mM ATP for 30 min at 30 °C. Accessibility of the BsiWI and StyI sites within the nucleosome was assayed by restriction digestion; the ClaI site, upstream of the promoter, was used to release the template after the remodeling reaction, followed by DNA purification and electrophoresis. (B) Immobilized +1 nucleosome remodeling assay was performed with or without TFIIH as in (A). (C) Cy3-labeled mononucleosome with a 77 bp linker DNA was treated with holo-TFIIH in the presence or absence of ATP for 30 min at 30°C. Reactions were quenched by addition of excess salmon sperm DNA, which strips TFIIH from the nucleosome. Samples were analyzed by non-denaturing polyacrylamide gel electrophoresis and scanned in the Cy3 channel. (D) Cy3-labeled mononucleosomes with 77 bp, 20 bp, or no linker DNA were remodeled with TFIIH as in (C). TFIIH-mediated nucleosome remodeling requires a long linker DNA.

### TFIIH remodels nucleosomes with long linker DNA

We next found that TFIIH has nucleosome remodeling activity on its own. Nucleosomes assembled on Cy3-labeled DNA were treated with TFIIH and ATP, TFIIH was removed by competition with excess DNA, and the products were analyzed by native polyacrylamide gel electrophoresis. Treatment of a nucleosome bearing a 77 bp linker (77N0) with 10-subunit holo-TFIIH converted it to naked DNA in an ATP-dependent manner (Figure 5C), indicating eviction of the histone octamer. TFIIH and SWI/SNF showed comparable remodeling activity at 200 nM (Figure 5C). Titration indicated that TFIIH-mediated remodeling saturates at approximately 100 nM or below (Figure 5D: 77N0).

The intrinsic nucleosome remodeling activity of TFIIH is attributable to the Ssl2 DNA translocase. TFIIH lacking Ssl2, purified as previously described (Murakami et al. 2012), had no nucleosome remodeling activity (Supplemental Fig. S7A), whereas core TFIIH, which lacks the Kin28 kinase, remodeled nucleosomes efficiently (Supplemental Fig. S7B). Remodeling therefore requires Ssl2 but not the kinase module, consistent with an earlier report that the other TFIIH ATPase, the Rad3 helicase, acts on single-stranded rather than duplex DNA *in vitro* (Sung et al. 1987).

In contrast to 77N0, treatment of 20N0 with TFIIH and ATP generated only a modest amount of naked DNA, and treatment of 0N0 gave almost undetectable product (Figure 5D). TFIIH therefore remodels nucleosomes with long linker DNA preferentially. We asked whether holo-TFIIH simply binds 77N0 more tightly than 20N0 or 0N0, and measured binding by electrophoretic mobility shift assay (EMSA) across a titration of TFIIH (Supplemental Fig. S7C). The affinity for 77N0 was indeed higher than for 20N0 and 0N0 (90%, 73%, and 63% bound at 100 nM TFIIH, respectively; Supplemental Fig. S7C), but this modest difference in binding does not account for the much larger difference in remodeling activity. The ATPase activities of Ssl2 and XPB increase with DNA length and plateau at ∼100 bp (Fishburn et al. 2015; Tomko et al. 2021), suggesting that the substrate preference for nucleosomes with long linker DNA is conferred by the Ssl2 ATPase itself. In budding yeast, nucleosomes are separated by ∼20 bp of linker DNA on average, the exceptions being the +1 and –1 nucleosomes, which flank long nucleosome-free regions. This length dependence may therefore help restrict remodeling by TFIIH, which otherwise binds naked DNA with high affinity, to promoter nucleosomes.

## DISCUSSION

This study makes three principal advances. First, we established an *in vitro* transcription system for the +1 nucleosome that recapitulates regulation observed *in vivo*: SAGA-dependent TBP loading, and a dependence on Mediator that increases when the +1 nucleosome constrains PIC assembly. Second, contrary to the prevailing model, we found that SAGA remains bound to the PIC after TBP loading and acetylates the +1 nucleosome in the context of the PIC. Third, we identified a previously unrecognized function of TFIIH in transcription initiation: remodeling of the +1 nucleosome. TFIIH, long known for its role in promoter opening, thus also couples transcription initiation to remodeling of the +1 nucleosome.

Upon stress, SAGA is recruited to the promoter by an activator (Figure 6, step 1). Because Pho4 is crucial for suppression of aberrant transcripts on the +1 nucleosome template but not on naked DNA (Figure 1C and Supplemental Fig. S3A), simultaneous interactions of SAGA with Pho4 and with the +1 nucleosome may orient SAGA such that TBP loading occurs only in the region surrounding the TATA element of the target gene (Figure 6, step 2). After TBP loading, SAGA remains bound to the promoter and is incorporated into the PIC. Because SAGA binds components of PIC-Med (Figure 3), promoter-bound SAGA may in turn facilitate PIC assembly by recruiting transcription proteins (Figure 6, step 3). Transient partial unwrapping of the +1 nucleosome may then allow productive engagement of the downstream PIC components with promoter DNA, and we propose that Mediator plays a crucial role in stabilizing promoter-bound TFIIH, thereby preventing rewrapping of the +1 nucleosome and subsequent occlusion of TFIIH from the promoter (Figure 6, step 4). Upon transcription initiation, TFIIH remodels the +1 nucleosome through the DNA translocase activity of Ssl2 (Figure 6, step 5). Cryo-EM studies of initially transcribing complexes (ITCs) have shown that TFIIH remains associated with ITCs in both yeast and human cells (He et al. 2016; Yang et al. 2022; Zhan et al. 2026), raising the possibility that TFIIH also facilitates early Pol II elongation through remodeling of the +1 nucleosome.

**Figure 6.**
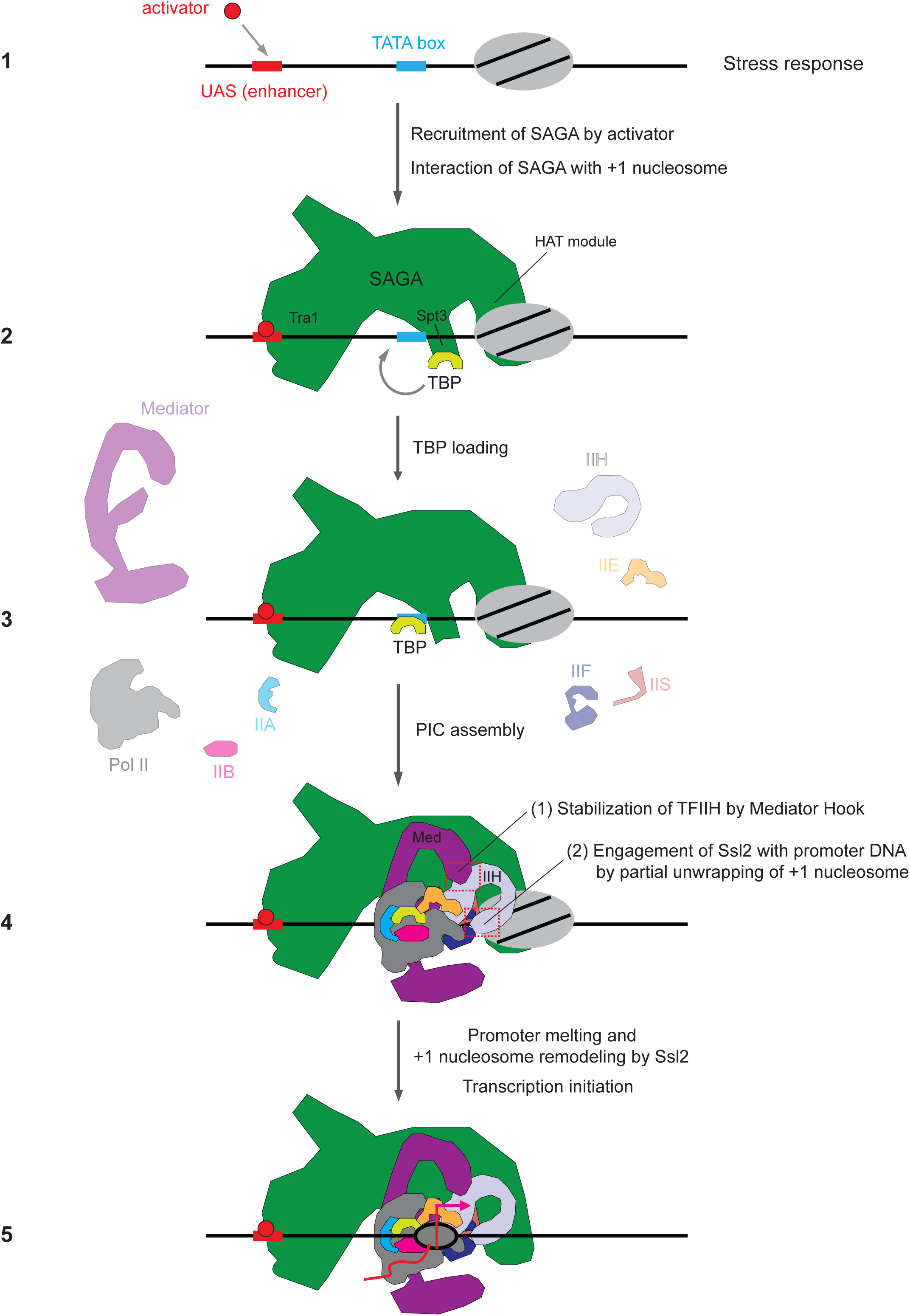
Cartoon model of +1 nucleosome transcription at SAGA-dependent promoters. Upon stress, an activator binds the upstream activating sequence (UAS) and recruits SAGA to the promoter (step 1). Engagement of SAGA with the activator and with the +1 nucleosome facilitates loading of TBP onto the TATA box of the target gene (step 2). SAGA remains bound to the promoter after TBP loading and may facilitate PIC assembly by recruiting transcription proteins (step 3), consistent with evidence that SAGA is required for Mediator recruitment *in vivo* (Bhaumik et al. 2004). When the +1 nucleosome constrains productive PIC assembly, transient partial unwrapping may allow Ssl2 to engage promoter DNA (step 4). Mediator is proposed to stabilize the resulting transcription-competent PIC, possibly by maintaining productive TFIIH– DNA engagement; the interaction between the Mediator hook and Rad3 of TFIIH is a candidate contributor to this stabilization (step 4). Ssl2 then drives promoter melting and remodeling of the +1 nucleosome, allowing transcription initiation (step 5).

The interaction of SAGA with PIC-Med is consistent with evidence obtained *in vivo*: artificial tethering of Mediator to the GAL1 UAS is sufficient to recruit SAGA in the absence of an activator (Lemieux and Gaudreau 2004), and at the same UAS activator-recruited SAGA is required for subsequent recruitment of Mediator (Bhaumik et al. 2004). We have also shown previously that short (53-residue) fragments of the SAGA core subunits Spt7 and Ada1 are each sufficient, when artificially tethered to a promoter, to recruit the transcription machinery and activate transcription (Sanborn et al. 2021). Because these fragments are too short to carry the Spt3/Spt8 TBP-delivery module or the HAT module, Spt7 and Ada1 must harbor sequences that recruit the transcription machinery independently of the canonical TBP-loading and histone acetylation functions of SAGA.

Addition of acetyl-CoA to the isolated PIC–Med–SAGA–nucleosome complex resulted in acetylation of histone H3 (Figure 4D), demonstrating that the SAGA HAT module engages the +1 nucleosome in the context of the PIC. As noted above, the human TFIIH subunit XPB (Ssl2 in yeast) interacts with KAT2A, the human Gcn5 homolog and catalytic subunit of the SAGA HAT module (Sandoz et al. 2019). Because the +1 nucleosome lies immediately adjacent to TFIIH (Chen et al. 2021a; Abril-Garrido et al. 2023; Wang et al. 2023), an analogous Gcn5–TFIIH interaction could position the HAT module close to the +1 nucleosome and thereby facilitate its acetylation. Our cryo-EM maps place SAGA-dependent density in contact with TFIIH and TFIIS (Supplemental Fig. S6), an arrangement compatible with such a mechanism, although the density is unassigned and a specific Gcn5–TFIIH contact remains to be tested directly.

The position dependence of Mediator stimulation indicates that a proximal +1 nucleosome exposes a requirement for Mediator that is largely masked on accessible DNA. Mediator may stabilize a productive PIC against the competing tendency of nucleosomal DNA to rewrap, thereby increasing the likelihood that transient DNA accessibility leads to successful initiation. Because a nucleosome is present on the less Mediator-dependent templates as well, the effect is more readily explained by an action on productive initiation than by a general requirement for Mediator during passage through the nucleosome. Since PIC assembly precedes the addition of NTPs and is followed by washing, our assay is sensitive to both the formation and the retention of competent complexes. Stabilization of productive TFIIH–DNA engagement is one possible molecular basis for the effect, but the present data do not distinguish this mechanism from a broader stabilization of the PIC (Supplemental Fig. S4).

Cryo-EM structures of the TFIID-bound PIC showed that TFIID binds the p8/p52 dimerization domain (Tfb2/Tfb5 in yeast) of TFIIH, and that TFIID and Mediator together sandwich and stabilize TFIIH (Chen et al. 2021b), suggesting that TFIID could contribute to productive PIC organization on nucleosomal templates through analogous stabilization of TFIIH. A previous *in vitro* study of human +1 nucleosome transcription using TFIID showed Mediator-stimulated transcription of the +1 nucleosome (Nock et al. 2012). In that study, the +1 nucleosome was installed 71 bp from the TATA box, a configuration that does not interfere with assembly of the TFIID-based PIC-Med (Chen et al. 2021b); accordingly, the authors proposed that Mediator facilitates early elongation (Nock et al. 2012). These findings leave open the possibility that Mediator supports different steps of transcription depending on promoter architecture, co-activator usage, and nucleosome position.

Unlike the ATPase domains of conventional chromatin remodelers, which engage the nucleosome at SHL–2 or SHL+2, the Ssl2 translocase does not bind the nucleosome core. Instead, Ssl2 binds linker DNA and uses the energy of ATP hydrolysis to peel DNA off the edge of the nucleosome, which we infer leads to histone eviction, since the remodeling product migrates as naked DNA (Figure 5C and 5D). This mechanism predicts eviction rather than sliding, and it accounts for the requirement for long linker DNA. During initiation, TFIIH draws downstream DNA toward the transcription start site and thereby evicts a promoter-proximal +1 nucleosome (Figure 5). The processivity of TFIIH is approximately 90 bp (Fazal et al. 2015), which may limit its ability to disassemble more distal nucleosomes; consistent with this limit, substantial amounts of stalled transcripts were observed on the distally positioned T77N template (Figure 4E). The limited processivity of TFIIH may therefore create a need for an additional remodeling activity at promoters where the +1 nucleosome lies farther downstream. We have recently shown that SWI/SNF is incorporated into the PIC when the +1 nucleosome is distally positioned (T81N) (Nagai et al. 2026), a situation in which TFIIH may be unable to reach the nucleosome efficiently. Such a switch between remodeling activities would support efficient initiation across promoters that differ in the position of the +1 nucleosome, and acetylation of the +1 nucleosome by SAGA may contribute by recruiting the bromodomain-containing SWI/SNF complex (Hassan et al. 2002).

There has been much discussion of why TFIIH is required for transcription of protein-coding genes in eukaryotes but has no counterpart in prokaryotes. Early studies proposed that eukaryotic genomes, unlike prokaryotic genomes, are not under negative superhelical tension, creating a requirement for TFIIH in promoter opening (Parvin and Sharp 1993). The so-called “built-in block” model instead proposes that XPB (Ssl2 in yeast) itself inhibits promoter opening, an inhibition relieved by its ATPase activity (Lin et al. 2005; Alekseev et al. 2017), thereby providing an additional point of regulation for class II transcription. A more recent cryo-EM study proposed that most Pol II promoters have evolved to impose less DNA distortion than those of bacterial polymerase or of eukaryotic Pol I and Pol III, rendering them TFIIH-dependent and hence more regulatable (Dienemann et al. 2019). Our results suggest a further, non-exclusive rationale: TFIIH may have evolved to meet the challenge posed by the chromatin environment, coupling promoter opening to removal of the +1 nucleosome.

## Supporting information

Nagai et al Supplemental Materials

## ACKNOWLEDGEMENT

We thank the Stanford Macromolecular Structure Group for technical support with the Prometheus Panta dynamic light scattering instrument (purchased with funding from the Stanford c-SHARP Program). Cryo-EM data collection was performed at the Stanford-SLAC Cryo-EM Center (S^2^C^2^) and Stanford cryo-electron microscopy center (cEMc). S^2^C^2^ is supported by the National Institute of General Medical Sciences (1R24GM154186). This research was supported by a gift from World Laureates Association to R.D.K.

## AUTHOR CONTRIBUTIONS

S.N. conceived the project and conducted experiments. S.N., D.-H.C., and R.H.M. performed cryo-EM analyses. H.Z. purified Mediator kinase module. S.N. and R.D.K wrote the manuscript with input from all authors.

## DECLARATION OF INTERESTS

The authors declare no competing financial interest.

## MATERIALS AND METHODS

### Purification of Pol II, GTFs, TFIIS, Mediator, Mediator kinase module, and SAGA

TFIIA, TFIIB, TBP, TFIIE, TFIIS, and Pho4 were available in recombinant form (Nagai et al. 2017; Nagai et al. 2026). Pol II, TFIIF, TFIIH, Mediator, and SAGA were isolated from *S. cerevisiae* as previously described (Nagai et al. 2017; Nagai et al. 2026). TFIIH lacking Ssl2 was purified from an *S. cerevisiae* strain harboring Tfb4-TAP, as previously described (Murakami et al. 2012). Protein purity was verified by SDS-PAGE and Coomassie brilliant blue staining (Supplemental Fig. S2). ΔCTD Pol II was prepared from a strain harboring a Protein A/TEV tag inserted in the Rpb1 subunit immediately preceding the consensus YSPTSPS repeats.

### Reconstitution of the +1 nucleosome

*Xenopus laevis* histones were expressed and purified from inclusion bodies as described (Dyer et al. 2004). For octamer preparation, lyophilized histones were resuspended in unfolding buffer (20 mM Tris-Cl pH 7.5, 7 M guanidine hydrochloride, 10 mM DTT) to a concentration of 1 mg/ml. Histones H2A, H2B, H3, and H4 were combined at a molar ratio of 1.1:1.1:1:1 and dialyzed three times against 1 L of refolding buffer (10 mM Tris-Cl pH 7.5, 2 M NaCl, 1 mM EDTA, 5 mM β-mercaptoethanol). The sample was concentrated and applied to a Superdex 200 10/300 size-exclusion column pre-equilibrated with refolding buffer. Peak fractions were pooled, concentrated, and flash-frozen in liquid nitrogen.

Nucleosomes were reconstituted as described (Dyer et al. 2004). A 160 bp nucleosome DNA fragment containing the transcription start sites and part of the 601 sequence was prepared by PCR using an upstream primer containing a BsaI site, which cleaves outside its recognition sequence. BsaI-digested DNA and histone octamer were mixed at a 1:1.1 molar ratio in the presence of 2 M NaCl and 10 mM DTT and incubated for 30 min on ice. The sample was transferred to a Slide-A-Lyzer 10K MWCO MINI device (Thermo Fisher) and salt-gradient dialyzed from 400 ml of 2 M KCl buffer against 2 L of 0.25 M KCl buffer over 30 h. Nucleosomes were then subjected to 5%–30% sucrose gradient centrifugation at 40,000 rpm for 13 h using an SW60 rotor. Peak fractions were flash-frozen in liquid nitrogen and stored at –80°C until promoter ligation.

Promoter fragments were prepared by PCR followed by BsaI digestion. Ligation of the promoter fragment (with a TEG-biotin tag at the upstream end) to the nucleosome core particle was performed in 50 mM Tris-Cl pH 7.5, 4 mM MgCl2, 1 mM ATP, 10 mM DTT, and 20 U/µl T4 DNA ligase; the molar amount of promoter DNA was kept below that of the nucleosome core particle. Ligation was performed at 4°C for 15 h. The resulting +1 nucleosome was then captured on streptavidin T1 Dynabeads (Thermo Fisher), washed several times with W-150 buffer (10 mM Tris-Cl pH 7.5, 150 mM NaCl, 1 mM EDTA), resuspended in TCA buffer (10 mM Tris-Cl pH 7.5, 1 mM DTT, 1 mM EDTA, 0.01% (w/v) NP-40), and stored at 4°C.

### *In vitro* transcription of the +1 nucleosome

Naked DNA templates were purified by treating the corresponding +1 nucleosome with histone-stripping buffer (10 mM Tris-Cl pH 7.5, 2 M NaCl, 1 mM EDTA). Purified Pol II, the six GTFs, TFIIS, Mediator, and SAGA (0.5 pmol each) were combined with nucleosome or DNA templates (0.5 pmol), and PIC formation was allowed to proceed for 30 min at room temperature (without ATP). After PIC assembly, beads were washed once with A-100 buffer (40 mM HEPES pH 7.5, 100 mM potassium acetate, 5 mM DTT, 5 mM magnesium acetate, 0.01 mg/ml BSA, 5% (v/v) glycerol), followed by addition of NTP mix (40 mM HEPES pH 7.5, 100 mM potassium acetate, 5 mM DTT, 5 mM magnesium acetate, 0.1 mg/ml BSA, 5% (v/v) glycerol, 0.01% (w/v) NP-40, 0.8 mM ATP, 0.8 mM GTP, 0.8 mM CTP, 20 µM UTP, 0.04 µM [α-32P]-UTP (2.5 µCi), RNaseOUT (5 U per reaction)) to initiate transcription. Transcription was allowed to proceed for 30 min at 30°C and quenched by adding 150 µl of stop buffer (10 mM Tris-Cl pH 7.5, 250 mM NaCl, 10 mM EDTA, 0.7% SDS, 0.013 mg/ml Proteinase K) for 30 min at 37°C. Reaction tubes were placed on a magnetic stand and the supernatant was recovered. Transcripts in the supernatant were ethanol-precipitated, resuspended in 8 µl of RNA loading dye (NEB), and run on a 6% denaturing PAGE gel as described.

### Immobilized +1 nucleosome remodeling assay

Binding reactions were set up as described for the transcription assay, scaled up 2-fold. After PIC assembly, beads were washed once with A-100 buffer, followed by addition of restriction enzyme mix (20 mM Tris-acetate pH 7.9, 50 mM potassium acetate, 10 mM magnesium acetate, 0.1 mg/ml BSA, 10–20 U of restriction enzyme). Where indicated, 1 mM ATP was included. Reactions proceeded for 30 min at 30°C (resuspended every 10 min) and were quenched by adding 150 µl of stop buffer for 30 min at 37°C. DNA was extracted by phenol/chloroform extraction, ethanol-precipitated, resuspended in 20 µl of 1× DNA loading dye, and run on a non-denaturing polyacrylamide gel (6% polyacrylamide, 0.5× TBE). The gel was stained with SYBR Green.

### Nucleosome remodeling assay (in solution)

Nucleosomes were prepared with Cy3-labeled yeast TEA1 promoter DNA and Xenopus histones. Remodeling reactions (20 µl) contained 25 nM nucleosome, 50–200 nM TFIIH, 20 mM HEPES pH 7.5, 100 mM potassium acetate, 5 mM DTT, 5 mM magnesium acetate, 0.1 mg/ml BSA, 5% (v/v) glycerol, and 1 mM ATP. Reactions proceeded for 30 min at 30°C and were quenched by adding 5 µl of stop buffer (20 mM HEPES pH 7.5, 1.65 mg/ml salmon sperm DNA, 6% (v/v) glycerol) for 5 min at room temperature. Electrophoresis was performed on a non-denaturing polyacrylamide gel (5% polyacrylamide, 0.2× TBE) for 60 min at 120 V at 4°C and scanned in the Cy3 channel.

### PIC recruitment assay

Binding reactions were set up as described for the transcription assay, scaled up 2-fold. After binding, beads were washed once with A-100 buffer and eluted in SDS-PAGE sample buffer. Following electrophoresis, membranes were immunoblotted with the specified antibodies.

### Reconstitution of the PIC-Mediator-SAGA-nucleosome complex

The PIC-Mediator-SAGA complex was assembled using a salt-dialysis protocol modified from (Nagai et al. 2026): 0.2 nmol of +1 nucleosome assembled on the *SNR20* promoter (or corresponding naked DNA) was mixed with 0.4 nmol TFIIE, 0.4 nmol TFIIS, 0.4 nmol TBP, 0.4 nmol TFIIA, 0.4 nmol TFIIB, 0.32 nmol TFIIF, and 0.2 nmol Pol II in buffer A-300 (40 mM HEPES-KOH pH 7.5, 300 mM potassium acetate, 5% glycerol, 2 mM magnesium acetate, 5 mM DTT, 0.15% 3-(decyldimethylammonio)propanesulfonate). After dialysis for several hours, 0.2 nmol TFIIH, 0.32 nmol TFIIK, 0.2 nmol Mediator, and 0.2 nmol SAGA were added, and the mixture was dialyzed overnight into buffer (50/40) (40 mM HEPES-KOH pH 7.5, 5% glycerol, 2 mM magnesium acetate, 5 mM DTT, 0.15% 3-(decyldimethylammonio)propanesulfonate, with the millimolar concentrations of potassium acetate/ammonium sulfate shown in parentheses). The mixture was then dialyzed into buffer (100/10) for 8 h and loaded onto a 10%–40% (v/v) glycerol gradient. Gradients were prepared with a Gradient Maker (BioComp) using two buffers differing only in glycerol content (10% and 40% (v/v); 40 mM HEPES-KOH pH 7.5, 100 mM potassium acetate, 2 mM magnesium acetate, 5 mM DTT). The complex was centrifuged at 60,000 rpm for 90 min (PIC-SAGA) or 70 min (PIC-Med-SAGA) in a Beckman SW60 Ti rotor and manually fractionated. For cryo-EM, samples were chemically stabilized by fixation in glycerol gradients containing increasing concentrations of glutaraldehyde (0%–0.1%, Electron Microscopy Sciences). Peak gradient fractions (200 µl/fraction) containing glutaraldehyde were quenched by addition of 100 mM Tris-Cl pH 7.5.

### EM specimen preparation

Peak fractions from glycerol gradient centrifugation were dialyzed overnight against buffer lacking glycerol (40 mM HEPES-KOH pH 7.5, 100 mM potassium acetate, 2 mM magnesium acetate, 5 mM DTT) and concentrated to approximately 1.2 mg/ml. Octyl-β-glucoside was added to a final concentration of 0.05% (w/v) before plunge-freezing with a Vitrobot Mark IV system (Thermo Fisher). 200-mesh Quantifoil R2/1 grids were glow-discharged for 40 s at 15 mA and 0.4 mBar using an EasiGlow glow discharger (Ted Pella, Inc.). A 3 µl aliquot of the PIC-Med-SAGA complex was applied to the grid, incubated for 10 s, blotted for 2 s, and plunged into liquid ethane at approximately 4°C and 98% chamber humidity. All grids were stored in liquid nitrogen prior to data collection.

### Cryo-EM imaging and data processing

The grid screening was performed on a 200-kV Glacios electron microscope (Thermo Fisher Scientific) and a preliminary dataset for most samples was collected with a dose rate of 10 e^-^/pixel/s and defocus range of −2.0 to −3.0 µm. The SerialEM package(Mastronarde 2005) was used for the automated data acquisition. The screening data was collected in the movie-mode images with 40 frames for each on a Gatan K3 direct electron detector camera (Gatan Inc) in counting mode with the magnification of 17,500× and the pixel size of 2.35 Å. The high-resolution movie-mode images were collected on a 300-kV Titan Krios electron microscope (Thermo Fisher Scientific) equipped with a K3 direct electron detector camera (Gatan Inc) in super-resolution mode and a BioQuantum energy filter (Gatan Inc) (20 eV in slit width) and with the exposure time of 4 seconds and the dosage per frame of 1.02 e/Å^2^/s and 40 total frames for each image and with the magnification of 64,000× and the pixel size of 0.7Å and the defocus range of −1.0 to −2.5 µm. All datasets were processed using Relion (Scheres 2012) and cryoSPARC (Punjani et al. 2017). MotionCor2 (Zheng et al. 2017) was used for the motion correction (with bin=2) after the importing of movie-mode images into Relion and CTFFIND4 (Rohou and Grigorieff 2015) was used for the estimation of the contrast transfer function (CTF).

### Dynamic light scattering (DLS)

PIC-Med-SAGA, with or without a +1 nucleosome, was assembled on the *SNR20* promoter and isolated by gradient centrifugation as described above. The isolated complex was dialyzed against DLS buffer (40 mM HEPES pH 7.5, 100 mM potassium acetate, 1 mM DTT, 2 mM magnesium acetate) for 12 h to remove glycerol. Forty consecutive DLS measurements were acquired for each capillary on a Prometheus Panta instrument (NanoTemper Technologies) at 75% LED power, 100% laser power, and 25 °C. Technical duplicates of 10 µl of each sample were loaded into glass capillaries (Prometheus NT.48 Series nanoDSF Grade Standard Capillaries). Scattering curves and apparent hydrodynamic radii were calculated with Prometheus Panta software, with solvent viscosity estimated from the composition of the DLS buffer.

## Notes

### Competing Interest Statement

The authors have declared no competing interest.

