## Supplementary material for "Reconstitution of +1 nucleosome transcription reveals coordinated functions of SAGA, Mediator, and TFIIH": Nagai et al Supplemental Materials

A

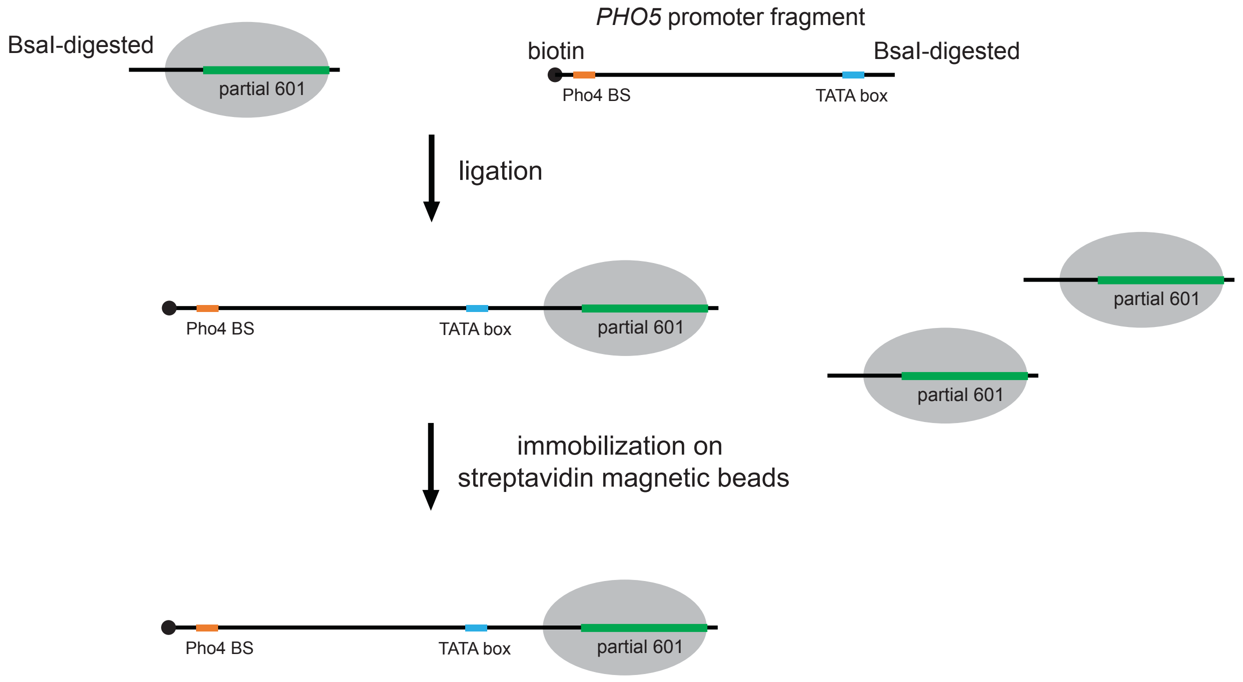


B

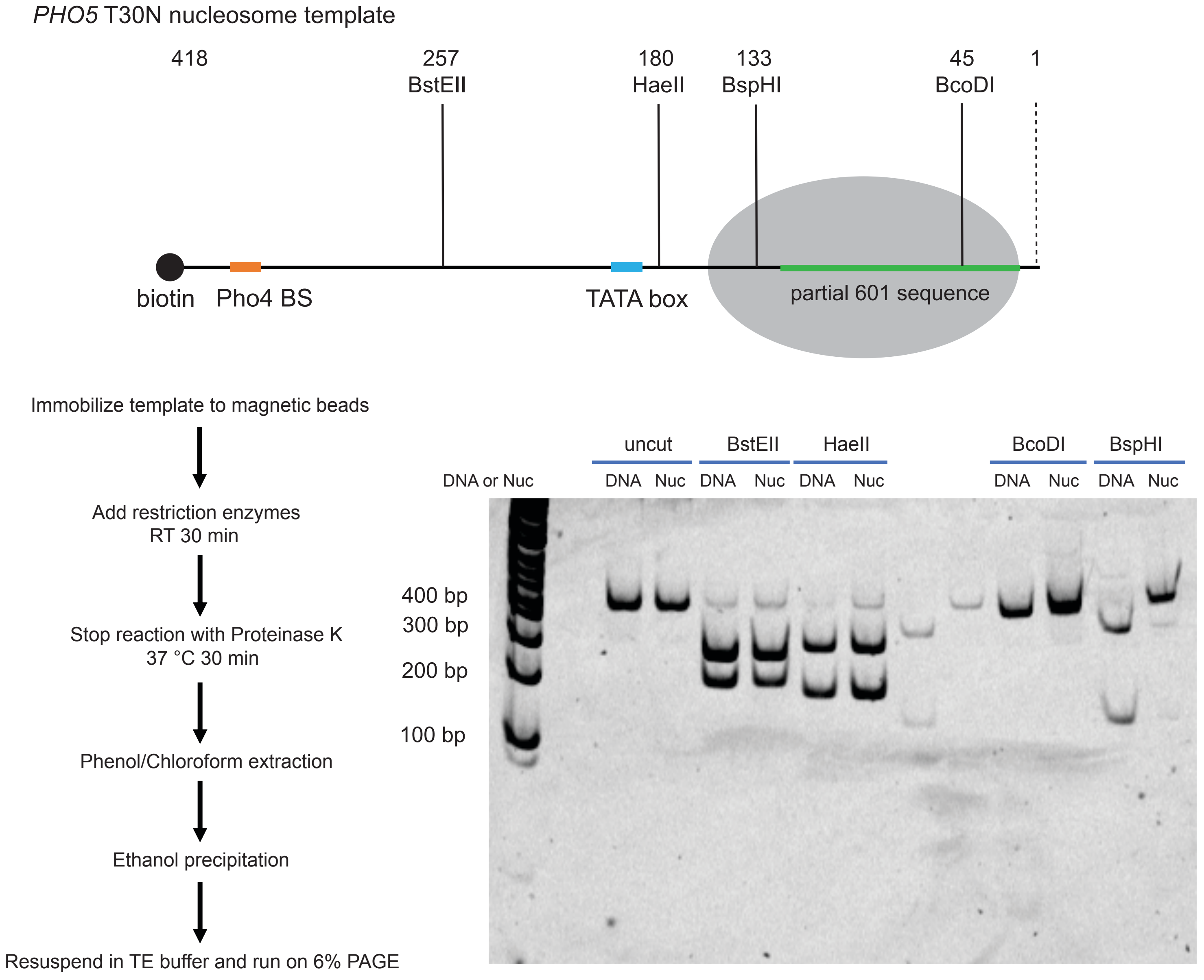


Supplemental Fig. S1

**Supplemental Figure S1. Construction and validation of the +1 nucleosome template.**

(A) Strategy for site-specific installation of the +1 nucleosome. A 160 bp fragment carrying the transcription start sites and the upstream part of the 601 positioning sequence was digested with the type IIS enzyme BsaI, which cleaves outside its recognition sequence and therefore leaves a defined overhang without introducing extraneous sequence, and was assembled into a mononucleosome. The mononucleosome was ligated to a BsaI-digested *PHO5* promoter fragment bearing a TEG-biotin tag at its upstream end, and ligation products were immobilized on streptavidin magnetic beads. Because only the promoter fragment carries the biotin tag, unligated nucleosomes and nucleosomes ligated at the incorrect end are removed during immobilization.

(B) Restriction endonuclease accessibility mapping of the *PHO5* T30N nucleosome template. (Top) Positions of the BstEII, HaeII, BspHI, and BcoDI sites relative to the biotinylated upstream end of the 418 bp template; the BspHI and BcoDI sites lie within the nucleosome and the BstEII and HaeII sites lie in the upstream promoter region. (Bottom left) Workflow. Immobilized templates were digested for 30 min at room temperature, and the reactions were stopped with Proteinase K at 37 °C for 30 min, followed by phenol/chloroform extraction, ethanol precipitation, and electrophoresis on a 6% polyacrylamide gel. (Bottom right) Naked DNA (DNA) and nucleosomal (Nuc) templates were digested with the indicated enzymes. Sites within the promoter region were cleaved on both templates, whereas the BspHI and BcoDI sites were cleaved on naked DNA but protected on the nucleosomal template, confirming that the histone octamer occupies the intended position.

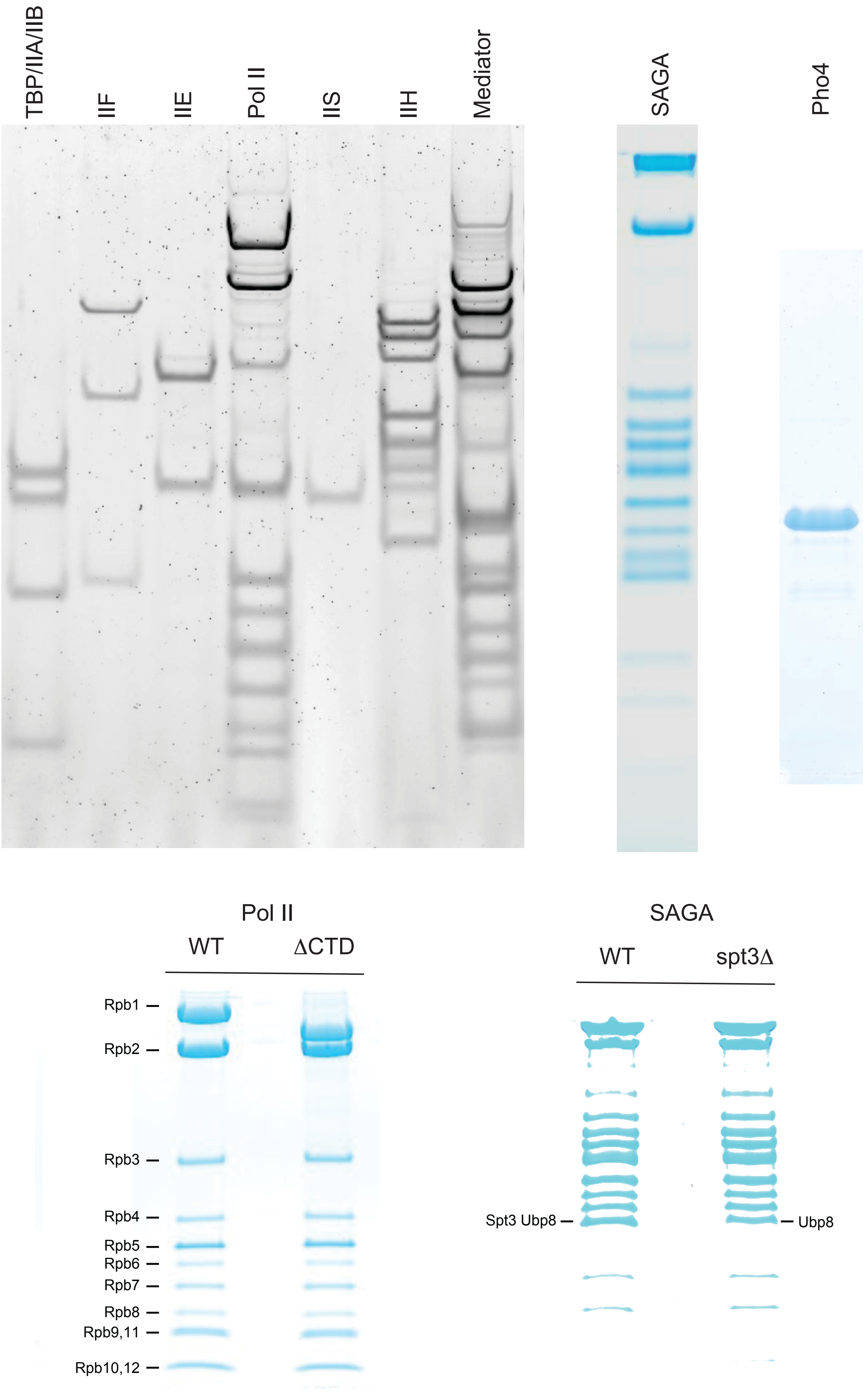


Supplemental Fig. S2

**Supplemental Figure S2. Purified proteins used in this study.**

(Top) SDS-PAGE of the purified transcription proteins, stained with Coomassie brilliant blue: TBP/TFIIA/TFIIB, TFIIF, TFIIE, Pol II, TFIIS, TFIIH, Mediator, SAGA, and Pho4. (Bottom left) Wild-type Pol II and ΔCTD Pol II. The twelve subunits are indicated; Rpb1 migrates faster in the ΔCTD preparation, consistent with removal of the CTD repeats, and the remaining subunits are present in unaltered stoichiometry. (Bottom right) Wild-type and *spt3*Δ SAGA. The Spt3 band is absent from the *spt3*Δ preparation, which otherwise retains the full complement of SAGA subunits.

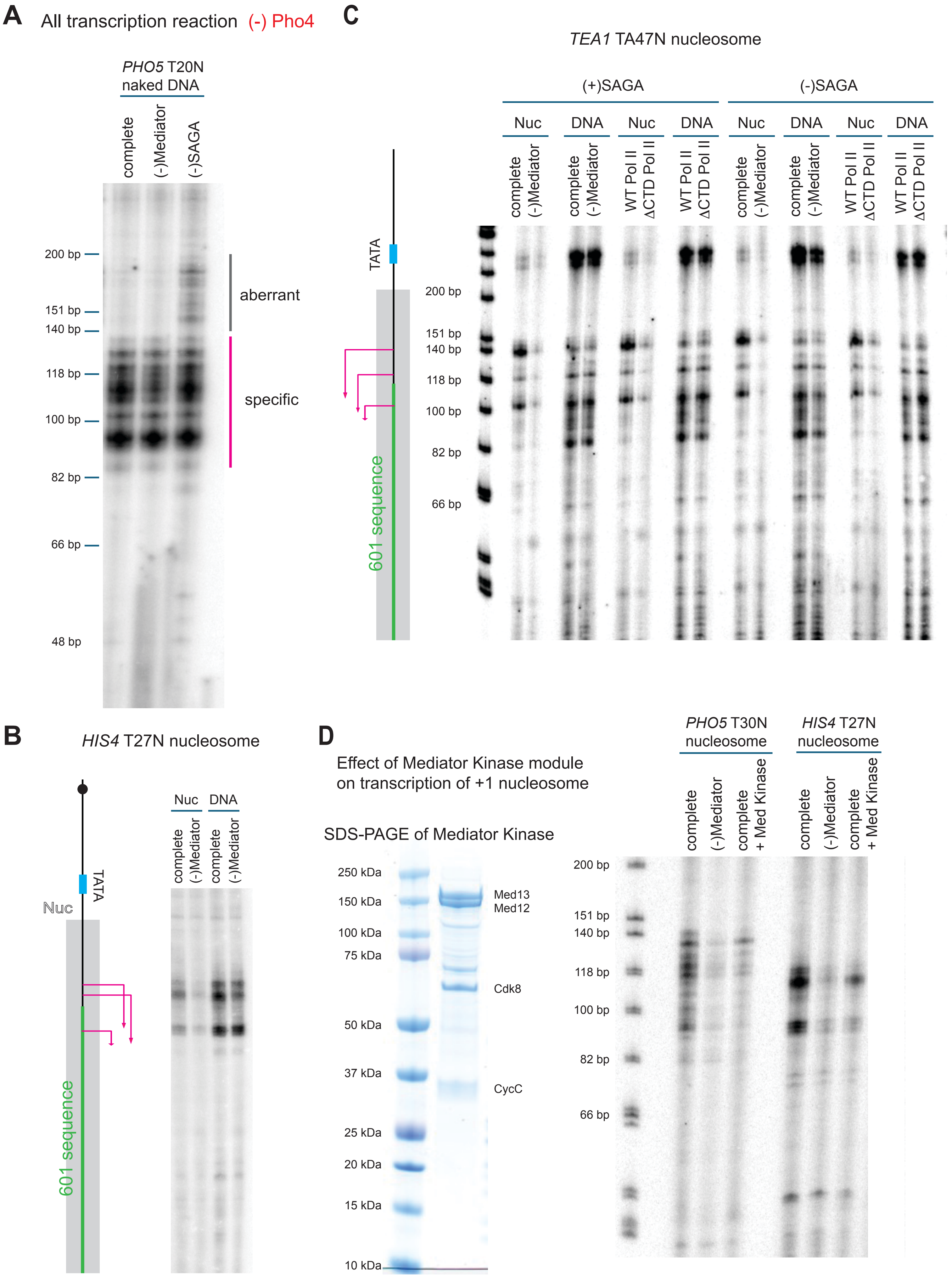


Supplemental Fig. S3

**Supplemental Figure S3. Additional transcription reactions on naked DNA and on the *HIS4* and *TEA1* +1 nucleosomes.**

(A) Transcription of the *PHO5* T20N naked DNA template in the absence of Pho4, with the complete set of factors, without Mediator, or without SAGA. Aberrant transcripts appear when SAGA is omitted, but not when Mediator is omitted, showing that SAGA suppresses aberrant initiation on naked DNA without a requirement for Pho4. Specific and aberrant transcripts are indicated by magenta and grey bars, respectively.

(B) Transcription of the *HIS4* T27N +1 nucleosome (Nuc) and the corresponding naked DNA (DNA), with the complete set of factors or without Mediator. Positions of the TATA box, the 601 sequence, and the transcription start sites (magenta arrows) are shown in the cartoon at left. Mediator strongly stimulates transcription of the nucleosomal template but has little effect on naked DNA.

(C) Transcription of the *TEA1* T47N +1 nucleosome and the corresponding naked DNA in the presence [(+)SAGA] or absence [(–)SAGA] of SAGA, with the complete set of factors, without Mediator, or with ΔCTD Pol II in place of wild-type Pol II. The requirements for Mediator and for the Pol II CTD on the nucleosomal template persist without SAGA, indicating that neither dependence is mediated by SAGA.

(D) Effect of the Mediator kinase module. The four-subunit kinase module, purified from a strain carrying Med12-TAP, was added to complete transcription reactions on the *PHO5* T30N and *HIS4* T27N +1 nucleosome templates. Reactions without Mediator are shown for comparison. Addition of the kinase module reduced transcription to approximately the level observed in the absence of Mediator.

### Immobilized PIC assembly assay

Immobilize *PHO5* T30N nucleosome to magnetic beads

↓

Add Pol II, GTFs, TFIIS, Med, SAGA, Pho4
PIC assembly RT 30 min

↓

Take off supernatant (S fraction)

↓

Wash unbound proteins (W fraction)

↓

Resuspend beads in SDS-loading buffer (B fraction)

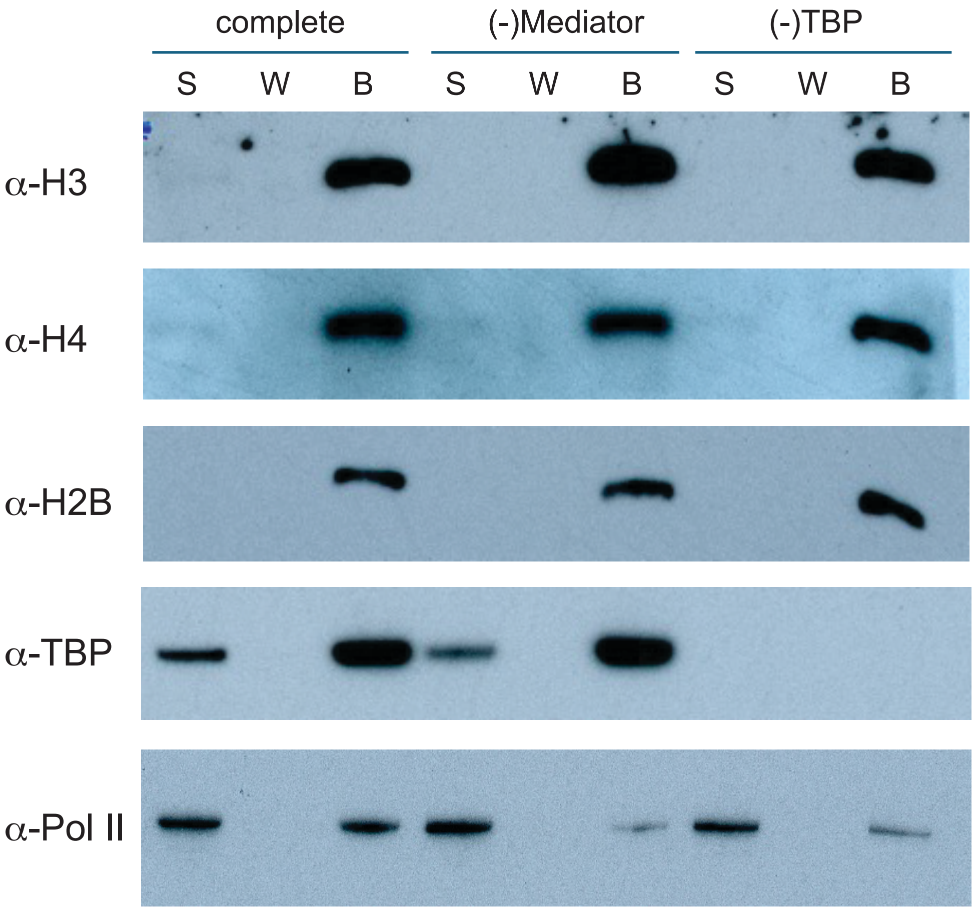


Supplemental Fig. S4

**Supplemental Figure S4. PIC assembly on the immobilized *PHO5* T30N +1 nucleosome.**

(Top) Workflow. Pol II, the GTFs, TFIIS, Mediator, SAGA, and Pho4 were incubated with the immobilized *PHO5* T30N nucleosome for 30 min at room temperature. The supernatant (S), a subsequent wash (W), and the material remaining on the beads (B) were analyzed. (Bottom) Immunoblots of the three fractions from the complete reaction, a reaction without Mediator, and a reaction without TBP, probed with antibodies against histones H3, H4, and H2B, TBP, and Pol II. Histones were recovered quantitatively in the bead fraction in every case, confirming equal template loading. TBP was retained on the template independently of Mediator, whereas retention of Pol II was strongly reduced when Mediator was omitted and was minimal when TBP was omitted, indicating that Mediator facilitates PIC assembly on the immobilized *PHO5* T30N nucleosome.

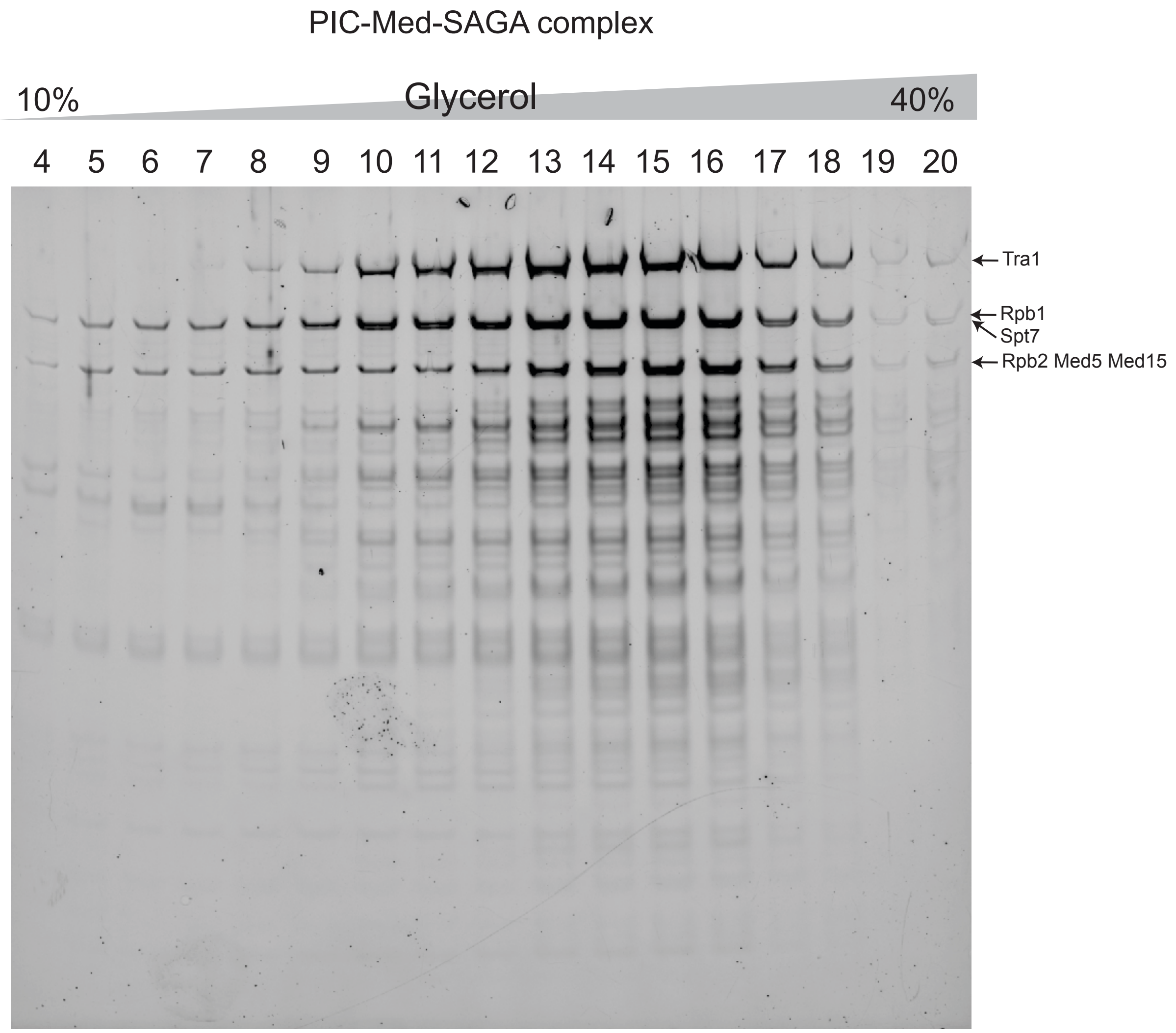


Supplemental Fig. S5

**Supplemental Figure S5. Isolation of the PIC-Med-SAGA complex by glycerol gradient sedimentation.**

SDS-PAGE of fractions of a 10%–40% glycerol gradient of PIC-Med-SAGA assembled on *SNR20* promoter DNA. Positions of Tra1 (SAGA), Rpb1 and Rpb2 (Pol II), Spt7 (SAGA), and Med5 and Med15 (Mediator) are indicated. Subunits of all three complexes co-migrate with a single peak centered on fractions 14–16, consistent with formation of a stoichiometric PIC-Med-SAGA assembly rather than independent sedimentation of its components.

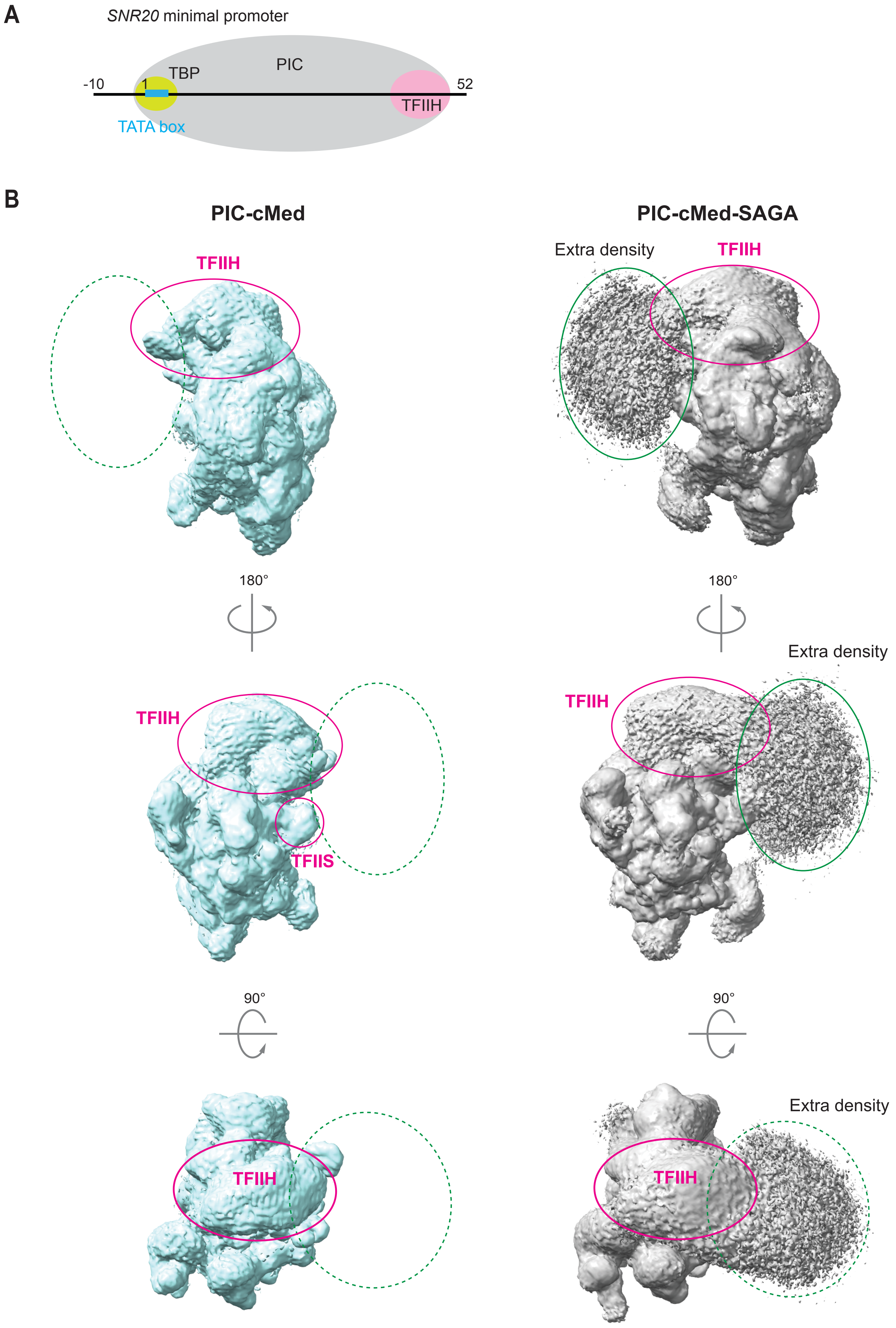


Supplemental Fig. S6

**Supplemental Figure S6. Cryo-EM reconstructions of PIC-cMed and PIC-cMed-SAGA.**

(A) The *SNR20* minimal promoter fragment used for assembly, extending from –10 to +52 relative to the 5’-edge of the TATA box, with TBP, and the approximate footprints of the PIC and TFIIH indicated. The fragment is too short to accommodate SAGA bound at a site distal to the PIC.

(B) Reconstructions of PIC-cMed (left, cyan) and PIC-cMed-SAGA (right, grey), shown in three orientations. SAGA-dependent extra density (green outline) contacts TFIIH and TFIIS in PIC-cMed-SAGA and is absent from the corresponding region of the SAGA-free control (dashed green outline). The extra density is too poorly resolved to permit assignment of individual SAGA subunits.

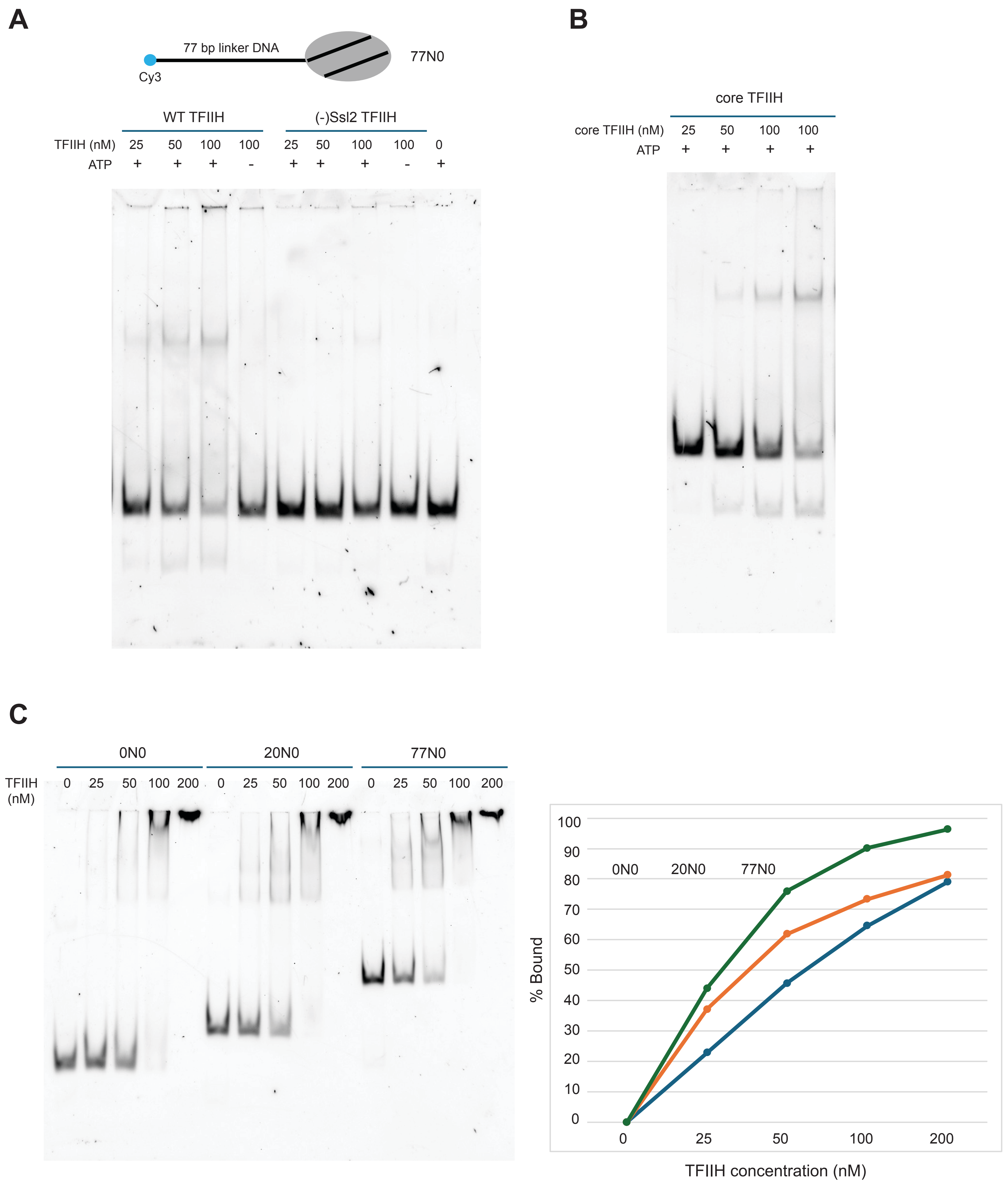


Supplemental Fig. S7

**Supplemental Figure S7. Requirements and nucleosome preference of TFIIH remodeling activity.**

(A) A Cy3-labeled nucleosome with a 77 bp linker (77N0; cartoon above) was incubated with wild-type TFIIH or with TFIIH lacking Ssl2 [(–)Ssl2 TFIIH] at the indicated concentrations, with or without 1 mM ATP, for 30 min at 30 °C. Reactions were stopped with excess salmon sperm DNA and analyzed on a native polyacrylamide gel scanned in the Cy3 channel. Wild-type TFIIH converted the nucleosome to free DNA in an ATP-dependent manner, whereas TFIIH lacking Ssl2 was inactive, identifying Ssl2 as the subunit required for remodeling.

(B) As in (A), with core TFIIH, which lacks the Kin28 kinase module. Core TFIIH remodeled 77N0, showing that the kinase module is dispensable for remodeling.

(C) Binding of holo-TFIIH to nucleosomes with no linker (0N0), a 20 bp linker (20N0), or a 77 bp linker (77N0), measured by electrophoretic mobility shift assay in the absence of ATP. (Left) Native gel across a TFIIH titration. (Right) Quantitation of the fraction of nucleosome bound. TFIIH bound 77N0 somewhat more tightly than 20N0 or 0N0 (90%, 73%, and 63% bound at 100 nM TFIIH, respectively); this difference is smaller than the difference in remodeling activity among the three substrates (Figure 5D).

**Supplemental Video 1. Coordinated motions of Mediator and core TFIIH within the PIC.**

Three-dimensional variability analysis of the yeast PIC-Med cryo-EM data set, computed in cryoSPARC and displayed along the first variability component. Mediator and core TFIIH move in a correlated manner, consistent with structural coupling between the two complexes through the Mediator hook–Rad3 interface.
